# NMRQNet: a deep learning approach for automatic identification and quantification of metabolites using Nuclear Magnetic Resonance (NMR) in human plasma samples

**DOI:** 10.1101/2023.03.01.530642

**Authors:** Wanli Wang, Li-Hua Ma, Mirjana Maletic-Savatic, Zhandong Liu

## Abstract

Nuclear Magnetic Resonance is a powerful platform that reveals the metabolomics profiles within biofluids or tissues and contributes to personalized treatments in medical practice. However, data volume and complexity hinder the exploration of NMR spectra. Besides, the lack of fast and accurate computational tools that can handle the automatic identification and quantification of essential metabolites from NMR spectra also slows the wide application of these techniques in clinical. We present NMRQNet, a deep-learning-based pipeline for automatic identification and quantification of dominant metabolite candidates within human plasma samples. The estimated relative concentrations could be further applied in statistical analysis to extract the potential biomarkers. We evaluate our method on multiple plasma samples, including species from mice to humans, curated using three anticoagulants, covering healthy and patient conditions in neurological disorder disease, greatly expanding the metabolomics analytical space in plasma. NMRQNet accurately reconstructed the original spectra and obtained significantly better quantification results than the earlier computational methods. Besides, NMRQNet also proposed relevant metabolites biomarkers that could potentially explain the risk factors associated with the condition. NMRQNet, with improved prediction performance, highlights the limitations in the existing approaches and has shown strong application potential for future metabolomics disease studies using plasma samples.

## Introduction

Metabolomics, one of the latest ‘omics’ techniques, is a powerful, high-throughput analytical technique to characterize the small molecules involved in metabolic chemical reactions. The overarching goal of metabolomics is to assess metabolites qualitatively and quantitatively for their diagnostic, therapeutic, and prognostic potentials. Metabolomics analysis allows us to highlight the unique fingerprints that distinguish the normal and pathological states, helping in disease diagnosis and prognosis predictions. Important metabolomics biomarkers with proven diagnostic and therapeutic relevance in managing disease conditions have been detected in many areas, including Alzheimer’s disease, dementia, Parkinson’s disease, inborn errors of metabolism, diabetic retinopathy, and cardiovascular disease^1^. To date, nuclear magnetic resonance (NMR) and mass spectrometry (MS) are the two major technical platforms for generating metabolomics data^2^. These techniques routinely detect thousands of signals in diversified biological samples^3^. Metabolomic tools offer the advantages of being fast, cheap, and sensitive. However, with the continued increase in sophistication and the number of signals, it is still challenging in metabolomics data analysis^1,3,4^.

Our study focuses on NMR spectra analysis. Practical metabolite identification and quantification from NMR spectra suffer from several challenges^5^. First, NMR spectra are of high complexity, composing thousands of data points from hundreds of metabolites with diversified sensitivities for explorations^6^. It requires high computational power and dedicated features screening pipeline to fully extracted the metabolites informative signals. Second, clusters from different metabolites could overlap heavily, raising the challenge of accurate signal decompositions, especially for low-concentration metabolites within the neighbors of dominant metabolites^7^. Third, NMR spectra are sensitive to the interference of environmental factors (temperature, pH, etc.) and the instrument, which could result in signal pattern alterations and positional shifts compared to the reference library^8^. With all the challenges, currently, the most commonly used method for metabolite analysis within NMR spectra still heavily relies on the annotation from NMR experts. Using Chenomx, NMR experts manually map potential metabolite candidates within the reference library to the mixture and generate a final fitting report^9^. This analysis process is not only time-consuming and expert-dependent but could also be subjective, which may result in less reproducibility. The challenges and variations are responsible for the limited application and inefficient interpretation of NMR spectra, which hinder its utilization in clinical practice.

To overcome these challenges, several methods have been developed. Current efforts solving the automatic explorations of NMR spectra include correlation-based, Bayesian-based, and linear regression-based methods. Correlation-based methods are led by Statistical Total Correlation Spectroscopy (STOCSY) and its variations^10–14^. Due to the high-dimensional problem in the covariance matrix estimation and overlapping signals, these methods have yet to identify metabolites within NMR spectra accurately. For Bayesian-based methods, Batman customizes the library based on specialized knowledge first and uses Markov chain Monte Carlo (MCMC) to find the best alignment for each metabolite cluster within the library^15,16^. For linear regression-based methods, they assume the underlying mapping relationship between the mixture and the reference library is a linear-regression model. However, when fitting metabolites in the reference library as the predictors and estimating their quantitative contributions to the mixture, the performance could substantially differentiate from the ideal case owing to the unpredictable and inconsistent variations among NMR samples^17,18^. Besides, for other methods, rDolphin^19^ and AQuA^20^, requiring manual library adjustment or dedicated sample preparations to prefilter the uninterested components within the biofluid samples, need more application convenience^21^. Recently, in deep learning, many methods have also been proposed for NMR applications ranging from spectra noise processing to metabolites identification^22–24^. Despite all these efforts, the metabolite features accompanied by variations in the real NMR spectra are still less capturable. The automatic and accurate metabolite quantification remains an unsolved question.

Here we present NMRQNet, a robust metabolite identification and quantification method in NMR spectra. We took advantage of the computational power within the deep learning model: convolutional neural network (CNN)^25^ and gated recurrent unit (GRU)^26^. CNN, with achievements in image classification, highlights the spectral features that best separate different metabolites^27^. With the CNN-extracted signals, GRU is further applied to learn the positional dependencies for clusters from the same metabolite, given its capability in signal processing^28^. With the combination of two model architectures followed by an optimization pipeline, our proposed computational method can automatically and accurately quantify the dominant metabolites within plasma samples despite positional and pattern variations brought by environmental and internal factors. NMRQNet will no doubt fill the application gap between the metabolomics platform in the lab and clinical.

## Results

### Overview of NMRQNet

The goal of NMRQNet is to capture both pattern and positional features of metabolites and automatically quantify the metabolites from the plasma samples (Fig. 1a). The first step, CRNN in NMRQNet, is a deep learning model consisting of two modalities. The first modality is a CNN block composed mainly of convolutional and pooling layers. It aims to aggregate the signal features and extract a high-level representation of metabolite clusters within the mixture^25,29^. The second modality adopts a gated recurrent unit layer, which takes the summarized features along the chemical shift axis step by step into the recurrent node to memorize the positional dependencies among metabolite clusters^26^. Finally, several fully connected layers are embedded to summarize all the learned features and predict the probability of the presence or concentrations of different metabolites within the library. The architecture scheme of the CRNN model in NMRQNet is shown in Figure 1b.

**Fig. 1.**
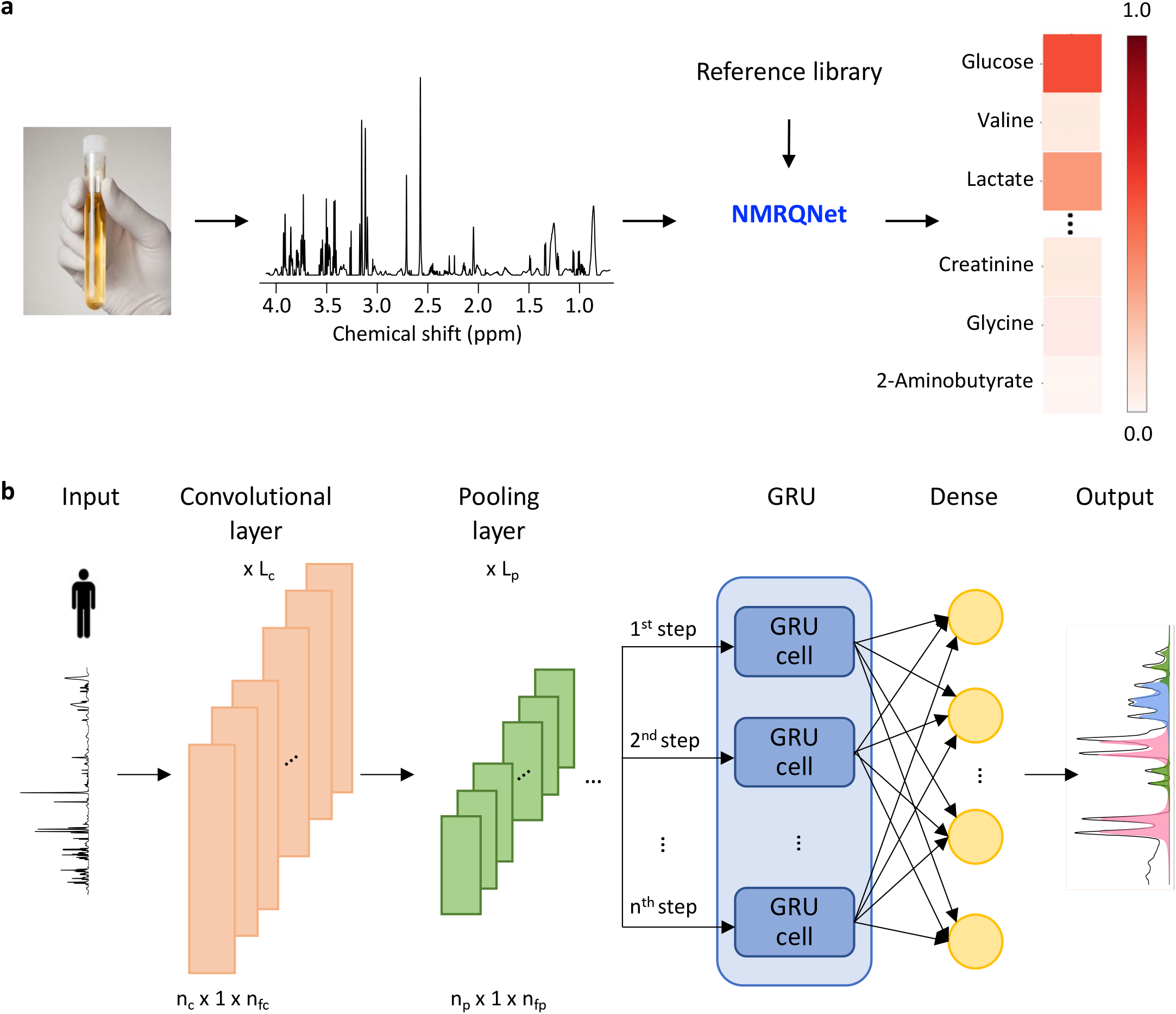
Overview of the NMRQNet workflow and CRNN model architecture. **a**, The plasma samples are collected to generate raw NMR spectra. Together with the reference library, NMRQNet automatically quantifies the relative concentrations for 38 metabolites within plasma samples. **b**, CRNN is a multi-model deep neural network consisting of two modalities for handling signal patterns and positional dependencies within the personal plasma NMR spectrum. The plasma NMR spectrum is input. The combination of multiple convolutional (L_c_) and pooling layers (L_p_) build up the convolutional blocks for signal pattern recognition. For each convolutional layer, the input and output sequences length is n_c_, and the number of filters is n_fc_. For each pooling layer, the length of the output sequence is n_p_, and the number of channels is n_fp._ The GRU cells are embedded for positional (chemical shift) memorization. The dense layer is applied to summarize the features and reconstruct the metabolites within the plasma based on the estimated concentrations.

Following that, for the second part of NMRQNet, a stochastic hill climbing-based algorithm was applied to optimize the quantification performance further^30^. Stochastic hill climbing altered and monitored the metabolite-specific position and concentration optimization process to better align them with the corresponding clusters within the mixtures. After the optimization step, the residuals between the original and reconstructed spectrum were minimized further.

NMRQNet starts with CRNN to capture the complicated features within the mixture and provide the preliminary quantification results. The reference library’s position was globally corrected based on CRNN results. Finally, we used stochastic hill climbing to fine-tune metabolite relative concentrations. By combining the deep learning and optimization steps, NMRQNet can gradually explore the best quantification alignments for the metabolite candidates within the mixture.

### Simulation of the training set from the reference library

With no public annotated NMR spectra database available, to address the lack of training samples for the CRNN model, we designed a simulation pipeline to synthesize annotated NMR spectra and embed variations from real spectra (Supplementary Fig. 1). We generated the whole NMR spectra for metabolomic candidates by applying Lorentzian-curve functions to the peak signatures (cluster, peak position, width, and height) within the reference library^31^. To overcome the discrepancy between the simulated and real NMR spectra, we implemented data augmentation techniques to include the noise and variations in each simulated spectrum^32^. In our experiments, we shifted each metabolite cluster within the neighboring region to simulate positional variations. We also scaled the peak widths to imitate pattern variations which are usually caused by environmental factors. Besides, signals from potential contamination and background noise were added to the simulation pipeline. Finally, the concentrations were randomly sampled from the fitted curve on manual concentration annotation results. This simulation pipeline generated sufficient training spectra to help CRNN learn the features of the real NMR spectra (Supplementary Fig. 1).

### CRNN accurately identifies and quantifies nine metabolites within known mixtures

To train a CRNN model, we simulated a data set from nine metabolites, including choline, glycine, myo inositol, cysteine, glutamate, glucose, lysine, tryptophan, and leucine. These nine metabolites represent a wide range of patterns and chemical shift values (Methods). We used ninety percent of the simulated samples for training and the rest for testing. We first trained the model to predict the probability of the presence of nine metabolites. After tens of epochs, the CRNN model converged on binary cross entropy loss function and had no further improvement in accuracy and F1-score.

To understand the performance of the CRNN model in accurate metabolite signal capturing, we first tested it on the simulated test data. In the predictions, CRNN can accurately predict the presence and absence of all nine metabolites for 92% of the test samples (Supplementary Table 1). Besides, CRNN can accurately identify at least six out of nine metabolites in all the samples. All the metabolites achieved an average accuracy of 0.978 to 0.995 (Supplementary Table 2). However, the question to be answered next is whether CRNN can still capture the right signals and be robust to the variations within the real NMR spectra.

To assess the performance of the CRNN model in handling positional variations within real experimental data, we physically mixed nine metabolites at fixed concentrations under three different temperatures (289*K*, 299*K*, 309*K*). We observed apparent positional variations among the spectra (Supplementary Fig. 2a). CRNN robustly identifies the presence and absence of most metabolites under positional variations. The prediction performance for choline and cysteine was slightly worse under 309*K*, which may result from the low concentrations and pattern alterations by baseline (Supplementary Fig. 2b).

**Fig. 2.**
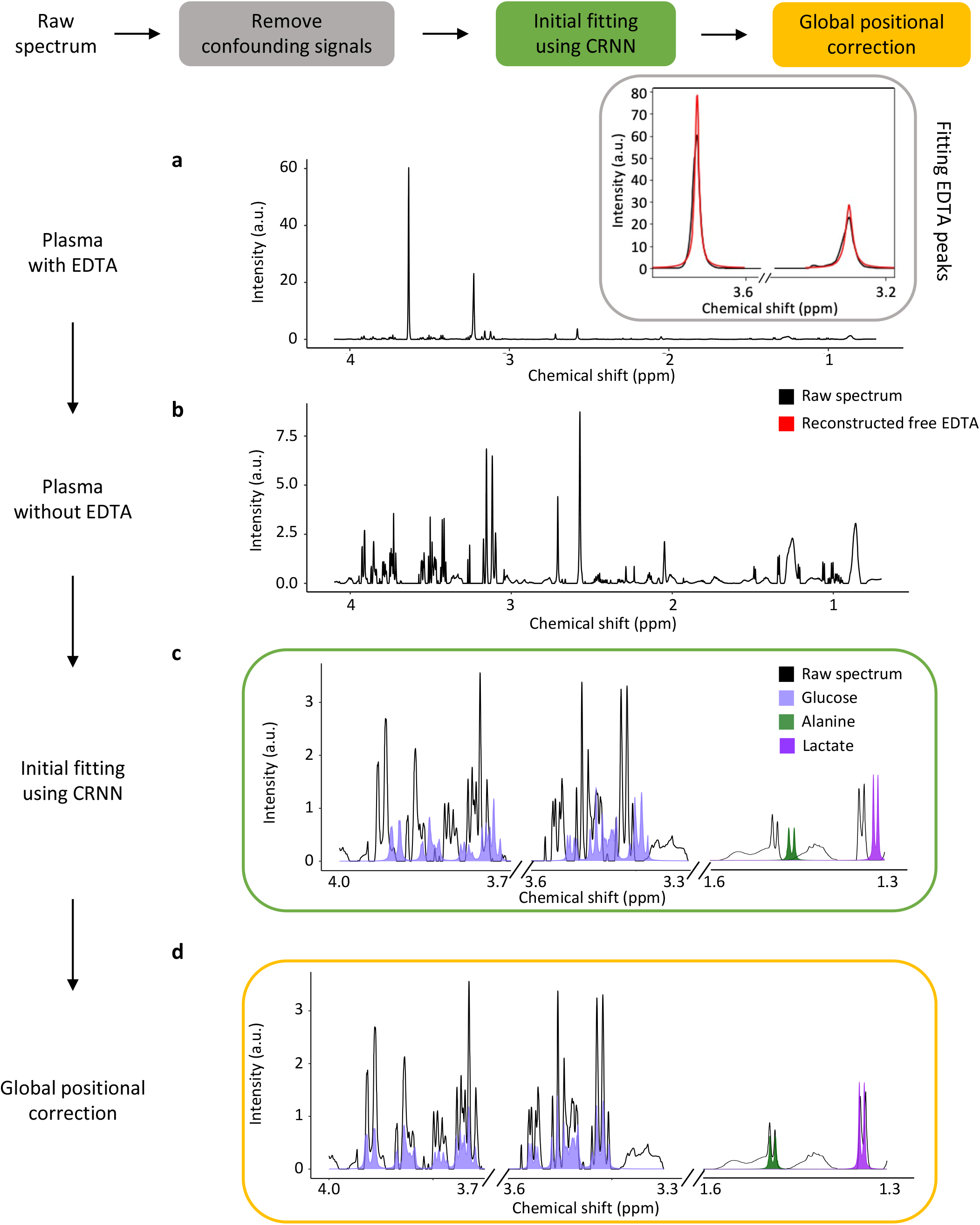
Initial CRNN fitting and global shift corrections for plasma NMR spectra. **a**, NMR spectrum for plasma with EDTA and the preprocessing process. Two free EDTA peaks have dominant concentrations within the mixture, suppressing metabolite signals. Stochastic hill climbing is applied to automatically target and fit the two free EDTA peaks and remove them from the plasma spectrum. **b**, Plasma NMR spectrum after removing the free EDTA peaks. Metabolite signals are clearer within the mixture. **c**, Initial fitting results from the CRNN model. CRNN predicted the concentrations for metabolites. The metabolite clusters are reconstructed based on the estimated concentrations and compared with the corresponding clusters within the plasma spectrum. The example shows the reconstruction of glucose (light purple), alanine (dark green), and lactate (purple). **d**, Correct the positions for all the metabolites globally. An optimal global shifting step is estimated with the shift that raised the highest local regional correlations. After the positional correction, all the metabolites are mapped to the neighboring regions of the corresponding metabolite clusters.

We further evaluate CRNN performance for quantifying metabolites within nine-metabolite mixtures. The quantification model for CRNN was trained on 10k simulated samples and predicted the area under the curve for different metabolites within the normalized mixture. For the simulated test data, CRNN can quantitively reconstruct all the metabolites correctly (Supplementary Fig. 3). With experimentally prepared NMR spectra, we applied CRNN to quantify potential metabolite candidates within the mixture. Based on the quantification results, we reconstructed the signals for different metabolites and mapped them back to the mixture spectrum (Supplementary Fig. 4). The mapping results show that most metabolites are accurately estimated, ranging from multiplet (glucose) to singlet (glycine). However, we still detected some imperfections within the prediction results. For example, our model may overestimate glutamate due to the low concentration, which makes it harder for the model to capture the actual metabolite signals mixed with the baseline. Besides, the impurities and background noise can lead to false-positive predictions. We implemented an optimization pipeline to overcome these challenges in later steps. Overall, our results indicate that the CRNN model has the potential to accurately target the metabolites signal mixed with variations and background noise.

**Fig. 3.**
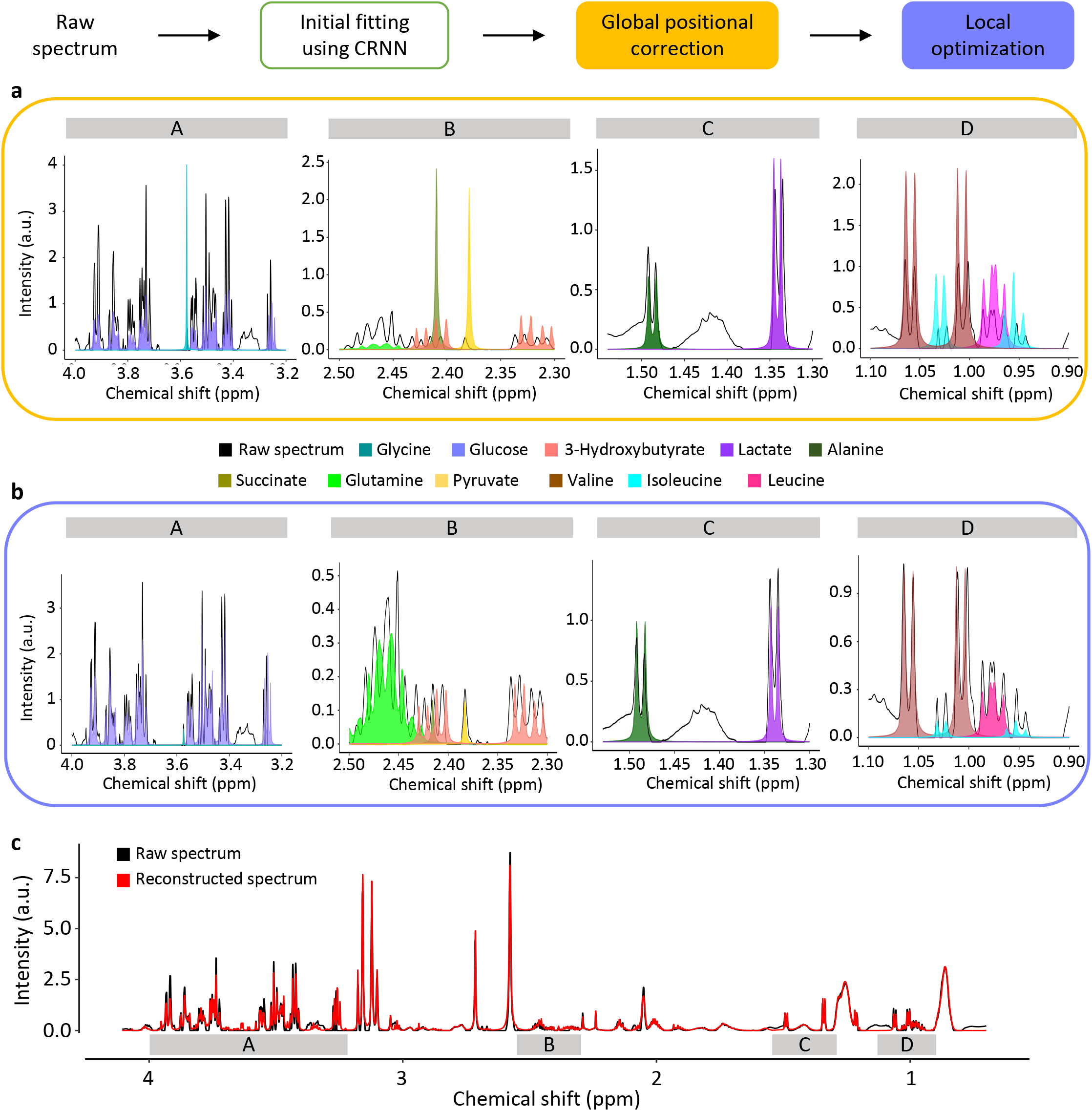
Local optimization with stochastic hill climbing. **a**, Metabolite reconstructions across different chemical shift regions after the global shift corrections. Alanine (dark green), lactate (purple), and 3-Hydroxybutyrate (coral) are well quantified. Succinate (olive green), pyruvate (yellow), glycine (turquoise), and metabolites in region D are overestimated. Glucose (light purple) and glutamine (green) are underestimated. **b**, Stochastic hill climbing is applied to optimize both the position and concentration for all the metabolites. After the local optimization, all the metabolites are better aligned with their targeted clusters within the plasma. **c**, Final mapping results from NMRQNet. The original raw plasma spectrum (black) overlaps well with the reconstructed spectrum (red) based on NMRQNet quantification results.

**Fig. 4.**
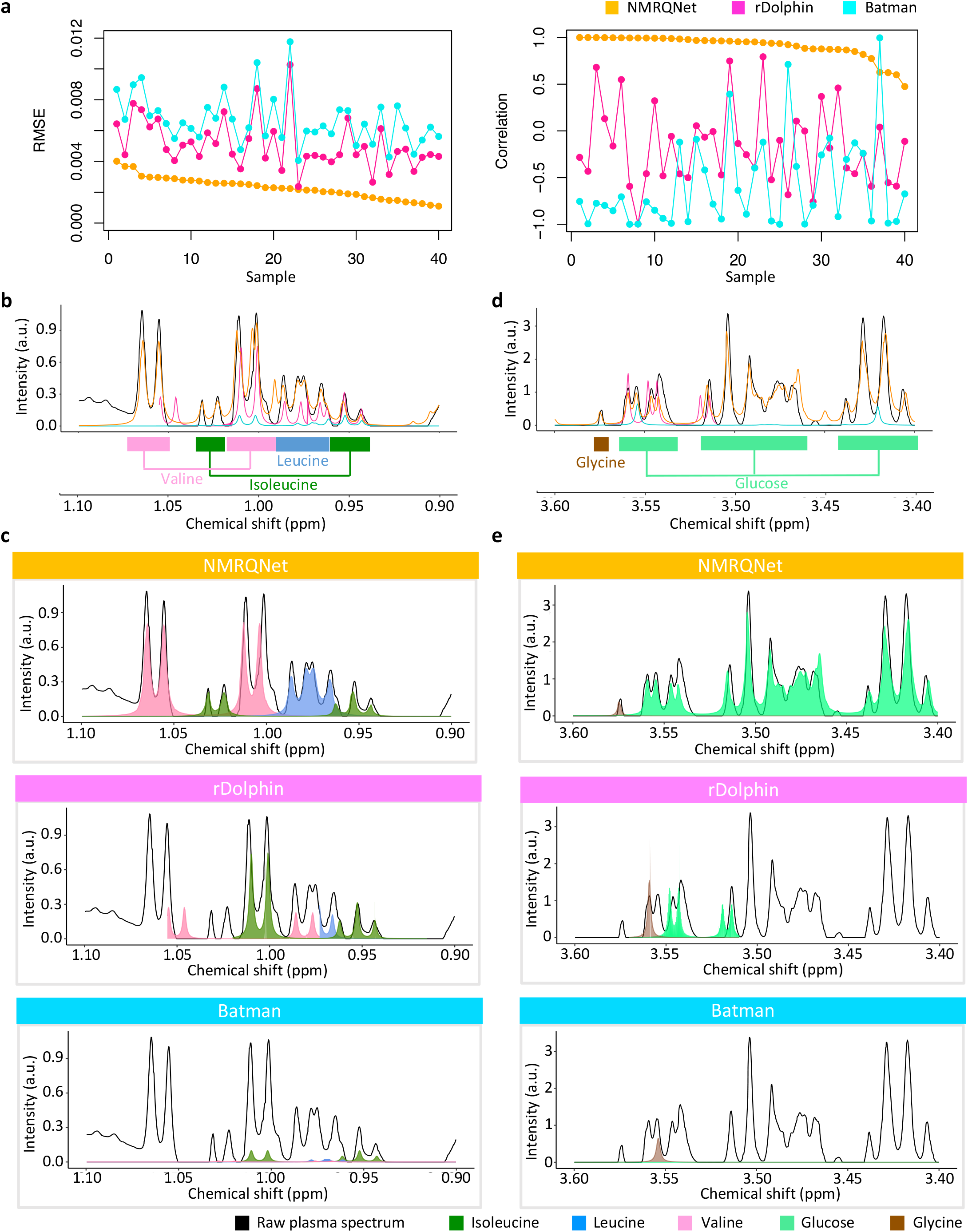
Performance evaluations across computational tools. **a**, Root mean squared error and correlation comparisons for metabolites in figure 4b among NMRQNet, rDolphin, and Batman across the 40 samples. From 0.9 to 1.1 ppm, areas under the curves for valine, isoleucine, and leucine in the original spectrum are calculated as ground truth. Sample-wise RMSE and correlation are calculated between the ground truth and the quantified areas under the curve based on the three metabolites. **b**, Metabolite annotations for clusters within the region of 0.9 to 1.1 ppm. Spectral reconstructions from different computational methods. **c**, Visualization of valine, isoleucine, and leucine reconstruction results across three methods. NMRQNet maps three metabolites to the right clusters with good quantification results. Both rDolphin and Batman have misaligned cases by mapping isoleucine signals to the valine clusters. **d**, Metabolite annotations for clusters within the 3.4 to 3.6 ppm region. Spectral reconstructions from different computational methods. **e**, Visualization of reconstruction results within 3.4 to 3.6 ppm across three methods. NMRQNet separates metabolites with different complexities (glycine and glucose) with good quantification performance. Both rDolphin and Batman can only reconstruct one cluster within the glucose and map the glycine singlet wrongly to the glucose cluster.

### NMRQNet precisely quantifies metabolites in plasma

Furthermore, to understand NMRQNet’s capability in dealing with the complexities within human biofluid samples, we applied NMRQNet to quantify metabolites within the plasma. For plasma analysis, we focus on the 39 commonly measurable metabolites as suggested by the IVDr (In Vitro Diagnostic Research) platform from Bruker. IVDr is developed to completely standardize and automate body fluids analysis for clinical and translational research^33^. We developed NMRQNet to achieve the same goal as IVDr but with open access. We specifically explored the signals within 0.7 to 4.1ppm, which has the most metabolite signals in plasma. Formate, with no peak within the selected region, is excluded from our plasma library. Thus, NMRQNet quantifies 38 metabolites in plasma samples. Besides, anticoagulants are always added within the blood tube, including EDTA, heparin, or Citrate, to prevent the blood from clotting^34,35^. We experimentally prepare the NMR spectra for the three anticoagulants. In addition, lipoprotein clusters are also major components within the plasma samples^36^. To capture lipoprotein signals, NMR expert located their positions within an external plasma spectrum, and we used normal distributions to fit those clusters (Methods). Finally, we take 24 normal distributions as the synthesized lipoprotein clusters. In total, we quantified 38 metabolites, 24 synthesized lipoprotein clusters, anticoagulants and DSS calibration standard.

To evaluate whether NMRQNet can automatically and accurately predict the metabolites within the plasma, we tested on 40 plasma samples from Mecp2 duplication syndrome (MDS) families with EDTA as the anticoagulant and generated the NMR spectra. Given that the free EDTA peaks have dominant concentrations which may obscure the signals from metabolites within the spectrum, we first applied stochastic hill climbing to target their positions and remove them from the spectrum (Fig. 2a)^30^.

As shown in Figure 2b, metabolite signals are more precise for further explorations with the removal of confounding signals. We next applied CRNN on the processed plasma spectra for initial fitting. Based on the quantification results, we reconstructed the metabolites within the mixture. For the reconstructed spectra, we found that metabolites in the sparse region reconstruct most of their corresponding clusters within the mixture (e.g., alanine and lactate). In contrast, metabolites within the overlapping regions can only reconstruct portions of the clusters (e.g., glucose) (Fig. 2c). To further improve the mapping performance, the prominent positional gaps between the plasma spectrum and the reference library are needed for corrections first.

To better align the metabolite clusters to the mixture, we optimize with global positional correction by shifting the reconstructed spectrum from initial fitting within the neighboring regions. From initial fitting results, CRNN has reconstructed most of the metabolite features within the NMR plasma spectrum. It makes it possible to figure out the optimal global shift step by calculating the regional correlations between every shifted reconstructed spectrum and the processed plasma spectrum. The global shifting step that gave the highest correlation across the regions will be used for global corrections for all the metabolite candidates. After the correction, we reconstructed the spectrum and observed that the reconstructed metabolites clusters were better aligned with their original clusters (Fig. 2d).

Although the fitting is improved, we still observed the under- or over-quantifications on some of the metabolites due to the overlapping or different sensitivities (Fig. 3a). To address these issues, we proposed a local optimization algorithm by shifting and scaling each metabolite individually and update using the stochastic hill-climbing algorithm ^30^. Specifically, in every step, each metabolite is finely tuned in concentration and position within the small regions. We only accepted the optimization if the mean square error decreased with the new concentration and position combination for the specific metabolite. Usually, within 500 iterations, we see a significant drop tendency, and the results converge in later iterations (Supplementary Fig. 5). After the local optimization, the reconstructed metabolite signals are better aligned with their corresponding clusters within the mixture (Fig. 3b).

**Fig. 5.**
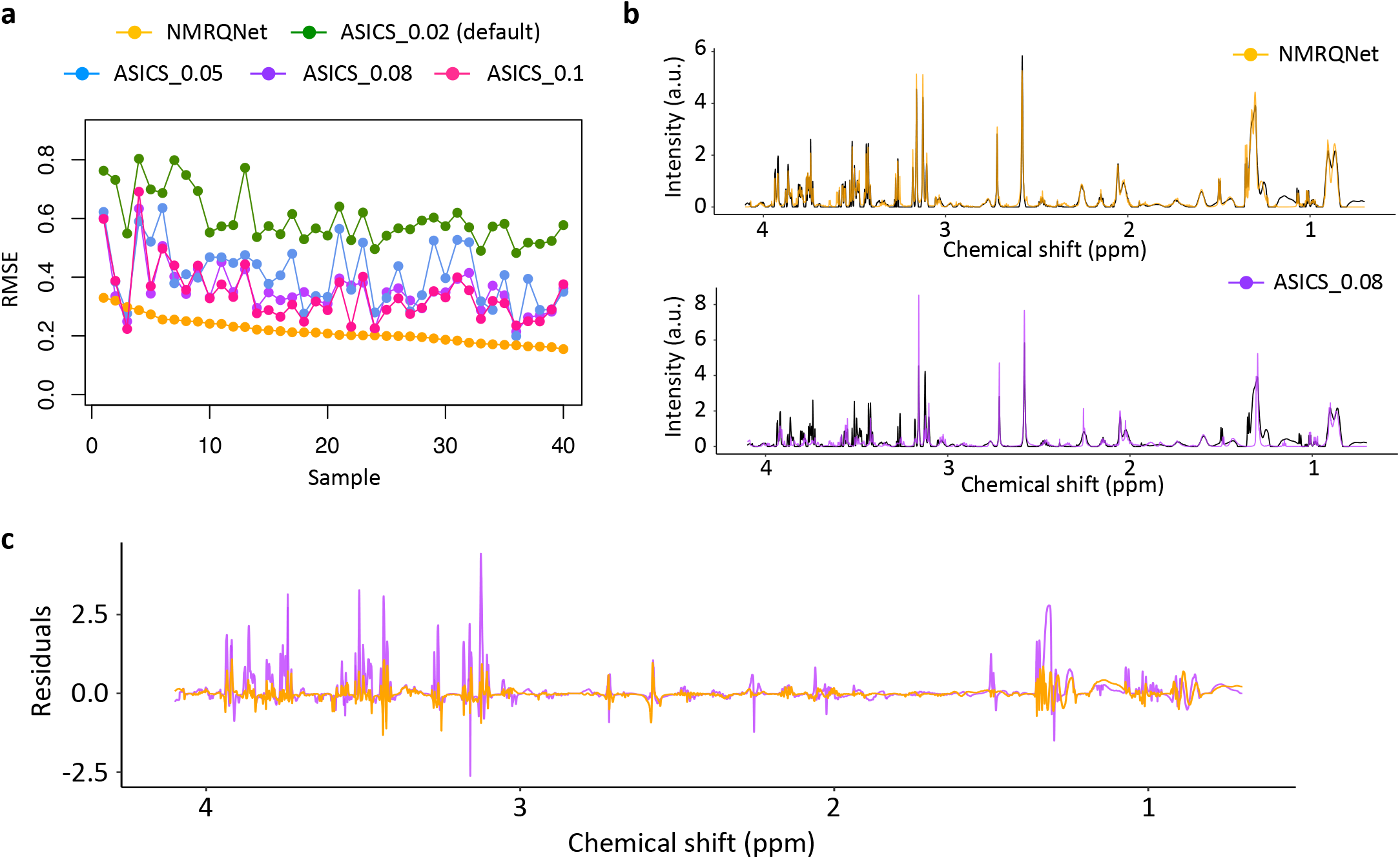
Performance comparison between NMRQNet and ASICS. **a**, Root mean square error (RMSE) between the raw plasma spectrum and the reconstructed spectrum from NMRQNet and ASICS. X-axis is the 40 plasma samples from Mecp2 duplication syndrome families. ASICS’s parameter max.shift is tuned from default 0.02 to 0.1. RMSE decreases as max.shift increases. ASICS’s results cluster above the NMRQNet results (orange). **b**, Reconstruction comparisons between NMRQNet (orange) and ASICS (purple) for the same sample. NMRQNet’s reconstruction spectrum overlaps well with the original spectrum. ASICS failed to reconstruct signals within 3 to 4 ppm in the original spectrum. **c**, NMRQNet and ASICS residual plots between the raw spectrum and the reconstruction spectra for the same sample in figure 5b.

Finally, based on all the fitting and optimization steps, we reconstructed the whole spectrum with the quantification results for all the potential components within the plasma (Fig. 3c). By aligning the reconstructed spectrum to the original one, the two spectra overlapped well and had small residuals across all the chemical shift regions. Besides the human plasma samples, plasma collected from mice and curated with different anticoagulants were also tested in Supplementary Fig. 6. Together, NMRQNet can precisely reconstruct the plasma spectrum with diversified anticoagulants from various species.

**Fig. 6.**
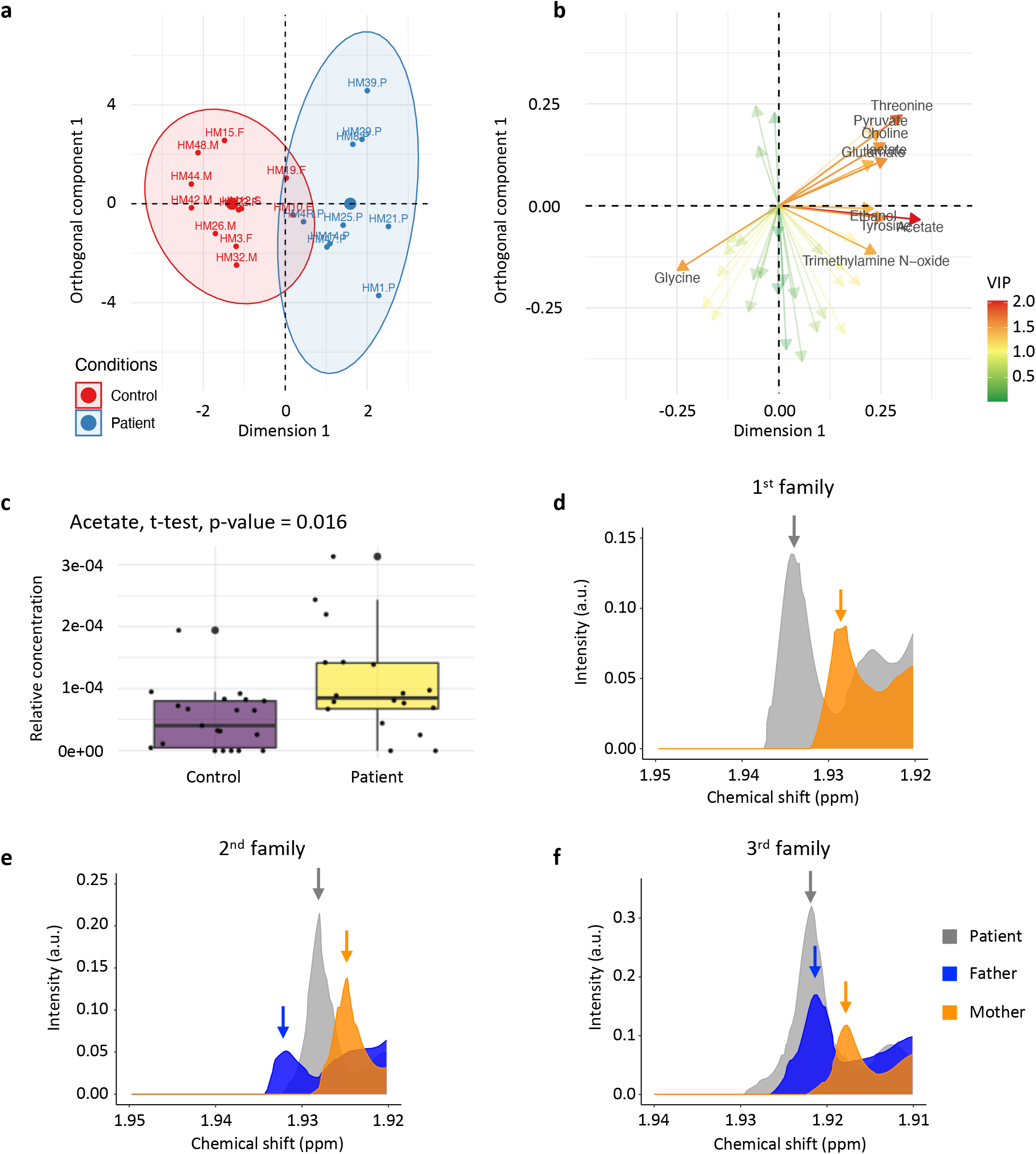
NMRQNet raises a novel metabolite biomarker within the downstream analysis. **a**, OPLS-DA plots on NMRQNet quantification results between the control and patient groups for the study of plasmatic metabolome within Mecp2 duplication syndrome families. **b**, Variable importance plot from the OPLS-DA analysis. **c**, Boxplot of the estimated acetate quantifications from NMRQNet between the control and patient group in the study of plasmatic metabolome within Mecp2 duplication syndrome families. T-test shows the significance of acetate with a p-value of 0.016. d, e, f, Acetate singlet within patients and parents’ plasma spectra in different families.

### NMRQNet outperforms other computational methods in metabolites quantification

To evaluate whether NMRQNet has improved quantification performance compared to the other state-of-art computational methods, we first compared NMRQNet with two approaches, which predict and quantitatively reconstruct metabolite-specific clusters within regions of interest. rDolphin uses a detailed region of interest (ROI) profile with chemical annotations in chemical shift regions, half bandwidth, and J-coupling. It further maximizes the line-shape fitting of signals in every analyzed ROI to explore the mapping cluster for every metabolite^19^. Batman also requires specified regions of interest and annotated chemical properties, including chemical shift positions and J-constant values. Moreover, it applied the Markov chain Monte Carlo (MCMC) algorithm to explore the metabolite quantification results automatically^15,16^. The methods were also tested on samples from Mecp2 duplication syndrome (MDS) families. For rDolphin, we applied their internal library for blood and kept the clusters within the region of 0.7 to 4.1 ppm. Across 40 plasma samples, rDolphin quantified 21 metabolites based on the region of interest. For Batman, with the ROI profiles for 757 metabolites, we extracted the 31 overlapping metabolites profiles between Batman and NMRQNet libraries. The primary metabolite quantification results among the three methods are compared in Figure 4.

The quantification performance was first evaluated by metabolite-specific area under the curve reconstructions (Fig. 4a). In the chemical shift region of Figure 4b, there are three major metabolites, valine, leucine, and isoleucine. We manually annotated each cluster and calculated the area under the curve as quantitative ground truth for each metabolite. Besides, the three methods (NMRQNet, rDolphin, Batman) all have metabolite-specific cluster reconstructions. Thus, they have the predicted area under the curve for each metabolite available. Each computational method was evaluated across three metabolites using two scorers: root mean squared error (RMSE) and Pearson correlation. Figure 4a shows the quantitative performance across 40 samples among three methods. NMRQNet is the best-performing method with the lowest RMSE and the highest correlation across most samples. rDolphin can provide reasonable quantification results in a few samples, while Batman failed in most cases. We further explored the reasons by visualizing the reconstructed clusters in Figure 4b-e.

We compared the methods on two regions of interest. As shown in Figure 4b-e, NMRQNet is overall the best performer by mapping all the metabolites to the right clusters within the mixture and estimating the concentrations accurately. In comparison, among the ROI profile-based methods^15,16,19^, rDolphin and Batman misaligned the isoleucine and glycine clusters to the neighboring clusters from other metabolites (Fig. 4c, e). This observation suggests the importance and convenience of automatic chemical shift adjustment during the mapping process. Relying heavily on manual positional adjustment in ROI profiles for both methods, they need to be more flexible in the automatic quantification of biofluid samples. Besides, rDolphin and Batman have limited power in dealing with high-complexity clusters. For glucose, they can only profile a cluster with two doublets. Discarding other dominant clusters from glucose, they are likely to lose its global picture and set the glucose quantification result subject to the influence of neighboring clusters. NMRQNet can overcome this limitation by having the whole spectrum for each metabolite included in the library.

ASICS is a method introduced more recently by ref. ^17,18^, which models the mixture spectrum as a linear combination of the library reference spectra. It adjusted the global shift for the reference library using R package speaq and further performed local distortions to minimize the residuals of local regression centered around each peak^37^. With the positional correction, the concentration for each component within the mixture is estimated from the linear regression model with the control of family-wise error rate (FWER). The library for ASICS is customizable. Thus, while testing ASICS, we used the same library as NMRQNet.

RMSE between the original plasma spectrum and the reconstructed spectrum from NMRQNet and ASICS is compared across 40 MDS-family plasma samples (Fig. 5a). The shift tolerance parameter in ASICS was tuned to optimize the mapping results^17^. In parameter explorations, we found no guaranteed performance improvement with increased shift tolerance (Supplementary Fig. 7). The optimal parameter is inconsistent among the closely generated samples, making it time-consuming to explore the parameter for every sample (Supplementary Fig. 8). Finally, we checked the results where ASICS performs better than NMRQNet. The higher residuals are raised from the lipoprotein clusters (Supplementary Fig. 9). Besides, in the spectral reconstruction comparison, metabolite signals within the 3 to 4 ppm region failed to be reconstructed by ASICS, which also led to more residuals, as shown in Figure 5b, c. Overall, based on the quantification results, NMRQNet consistently performed the best in regional explorations and whole-spectrum reconstruction.

### NMRQNet proposed a potential metabolite biomarker for Mecp2 duplication syndrome

To study how NMRQNet can help detect the potential metabolite biomarker across different conditions in disease, we collected plasma samples from Mecp2 duplication syndrome families with both patients and their healthy family members. Mecp2 duplication syndrome is a neurodevelopmental disorder disease that usually results from a heterozygous whole-gene duplication of MECP2. Patients with MDS are diagnosed with molecular genetic testing and may have intellectual disability, poor speech development, and other symptoms^38^. In our study, across 40 MDS family plasma samples, 19 samples are from patients, and the rest are from their healthy family members. From NMRQNet, we quantified the relative concentrations for 38 dominant metabolites by dividing the quantification results by the respective number of protons for different metabolites. Principal Component Analysis (PCA) was first applied to the estimated relative concentrations to screen potential outliers within the group (Supplementary Fig. 10). A patient sample is shown as an outlier, which could result from contamination in sample preparation. After excluding the outlier, we performed PCA (Supplementary Fig. 11) and Orthogonal Projections to Latent Structures Discriminant Analysis (OPLS-DA) on the rest of the samples^17,39^. In the OPLS-DA result, the control and patient groups are well separated, and the variables most important in separating the two groups were visualized in Figure 6a, b.

Further steps were taken to predict the control and patient group with logistic regression on 38 estimated metabolite concentrations. The performance based on the leave-one-out test is shown in Supplementary Fig. 12. Besides, we implemented a t-test between the groups to test the variable importance and significance. Acetate, with a p-value of 0.016, has significantly elevated concentrations within the patient group (Fig. 6c). To further confirm this potential biomarker in MDS, we specifically compared the acetate spectra peak between the patients and their parents, which offers reasonable genetic control. Across three families, we see the patients’ higher acetate peak intensities compared to their parents, which testify to our results (Fig. 6d-f). The relative concentrations estimated from NMRQNet capture the peak intensity elevation within NMR spectra across the groups. It supports the broad application of NMRQNet for plasma analysis in diversified disease areas to explore potential metabolite biomarkers.

## Discussion

NMRQNet is a novel approach for the automatic identification and quantification of metabolites within plasma NMR spectra. The method learns from metabolite-specific spectral patterns and captures cluster positional dependencies across the chemical shift axis. Furthermore, using regional correlation and stochastic hill climbing enables NMRQNet to automatically correct the variations and map the metabolite candidates to the fitted positions^30^. Also, our novel data augmentation-based simulation pipeline successfully embedded natural NMR features within the synthesized training set, which resolves the lack of annotated training data for NMR spectra (Supplementary Fig. 1)^32^. Hence, lower the entry bar and raise more possibilities for applying deep learning in computational metabolomics research.

Our experiments demonstrate that NMRQNet accurately reconstructs the whole NMR spectra with very low residuals compared to the original spectra, far more efficient than the traditional approach, and markedly better than previous computational-based methods that rely on Bayesian or linear-regression models (Figs. 4, 5)^15–18^. In particular, compared with ref. ^15,16,19^, which focused on the manual library adjustment and region of interest (ROI) driven quantification, the benefits of learning the positional correction parameter automatically and using the complete metabolite spectrum for mapping are clear. Different evaluation scores (RMSE, Pearson correlation, residuals) could be improved by NMRQNet, which also has better consistency among samples (Fig. 4a). Our results show the flexibility and robustness of deep learning-based model architectures for spectra processing. Remarkably, this was the case not only for quantifying metabolites within the plasma samples but also for automating the clinical assessment of plasma samples.

Our experiment also indicated that NMRQNet quantification results could be confidently applied to raise metabolite biomarkers, which could potentially explain the underlying disorders between the control and patient groups. However, further experiments were needed to confirm the potential biomarkers. Moreover, focusing on eliminating residuals between the original and reconstructed spectra, NMRQNet regularized less on the false-positive predictions, which requires final visualization check to filter metabolites with false-positive predictions. A better pre-filtering or post-processing step might be beneficial. Overall, the data augmentation implemented simulation pipeline overcame the data shortage issue for deep learning in the metabolomics area. The CRNN, with the combined feature-extraction power, also proves its capability in interpreting metabolite signals while processing spectral data. All these efforts could be transferred from the plasma area into other biofluid samples. Future efforts are needed to develop the biofluid-specific analysis pipeline and facilitate clinical diagnosis.

## Methods

### The CRNN model

We denote *X* = {*x*_1_, *x*_2_, *x*_3_, …, *x*_*m*_} as the input processed NMR spectrum for CRNN. Here, *m* is the length of the NMR spectrum. The input size for the nine-metabolite identification task is 10,000 dimensions, and for the nine-metabolite quantification problem, the input size is 40,000 dimensions. While for the plasma quantification model, the input size is 12,000 dimensions. After the input, a stack of convolutional layers acts as a metabolites’ signals scanner and extracts a feature panel X_f_ for each convolutional filter f. To prevent the model from overfitting, the dropout and batch normalization layers were added among the convolutional layers^40,41^. The dropout layers randomly assign zeros to network nodes. The batch normalization regularizes the mean and variance on each mini-batch from the output of the previous layer L and is formularized as,

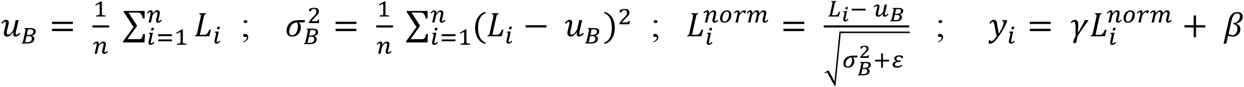

where *B* = [*L*_1_, *L*_2_, …, *L*_*n*_] is one mini-batch, n is the batch size, *u*_*B*_, 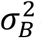 denotes the mean and variance of the mini-batch, 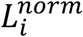 is the normalized version of *L*_*i*_, *γ* and *β* are the parameters used to scale and shift normalized values to get the output of batch normalization layer y_*i*_. A gated recurrent unit (GRU) layer with an update gate and a reset gate is employed to learn the positional dependency among metabolites’ clusters^26^, which is formularized as,

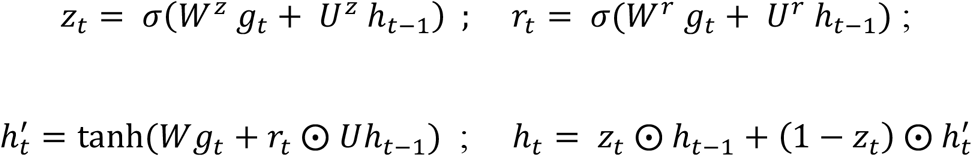

*z*_*t*_ and *r*_*t*_ are the update gate and reset gate for position step t, *σ* denotes the sigmoid function, *g*_*t*_ is the input at step t, *h*_*t−1*_ is the hidden state that holds the information from the previous t-1 steps, 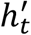 is the current memory content that uses the reset gate to store the relevant information from the past, *h*_*t*_ is the final hidden state for step t and will be passed down to the network, *W*^*z*^, *U*^*z*^, *W*^*r*^, *U*^*r*^, *W, U* are the weights learned during the training process. The output from GRU further connects the fully connected layers to generate the final output (Fig. 1b).

For the output, the final output layer for nine-metabolite identification is a nine-dimension vector, denoting the probability of the presence of nine metabolites. A binary cross-entropy loss is used to train this identification model. The output of the nine-metabolite quantification task is a ten-dimension vector, denoting the relative abundance for nine metabolites and DSS. We sequentially implemented two loss functions to optimize the weights of this deep-learning model. The first loss function was designed to minimize the quantitative gaps and the residuals between the original input and the reconstructed input, which is formulated as,

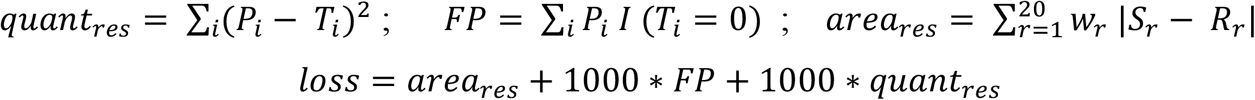

*P*_*i*_ is the predicted quantitative value for the i^th^ component, and *T*_*i*_ is the actual quantitative value for the i^th^ element in the library. *L* (.) denotes a boolean function with the value of 1 if *T*_*i*_ = *0*, but 0 for the rest of *T*_*i*_ values. Finally, r represents the regions evenly split from the spectra. *S*_*r*_ and *R*_*r*_ denote the sum of intensities in region r for the original spectrum, and the reconstructed spectrum, *w*_*r*_ represents the region-filtering parameter. The final loss function is the combination of all three terms trying to minimize the gaps between the original and reconstructed spectrum and regularize on the false-positive rate. 1000 is applied to scale the loss terms to a similar range. Once the model converged on the customized loss function after multi-epoch training, we further boosted the model performance by using mean square error as the loss function. Finally, for the deep-learning model designed for plasma analysis, the output is a 65-dimension vector, including dominant components within the plasma. Mean square error is used as the loss function to optimize the weights within the model.

### Training CRNN model

In the nine-metabolite identification task, during the training process, we evaluated the validation loss, accuracy, and f1-score at the end of each epoch to monitor the learning process. The weights for the convolutional, GRU and dense layers were initialized by Xavier normal distribution^42^. For quantification tasks (nine metabolites and plasma samples), the validation loss was monitored at the end of each epoch. He-normal initialization was used to initialize the weights in convolutional, GRU, and dense layers^43^. All the models were trained and optimized using the Adam algorithm^44^.

We simulated NMR spectra with known components and concentrations to train the model. For the nine-metabolite identification task, 70k annotated spectra were synthesized. Besides, we simulated 10k spectra for the nine-metabolite quantification model and 20k spectra for the plasma quantification study. To train and perform a thorough evaluation of these models, we left 10% of all the simulated datasets as the test set. For the rest of the samples, 90% of the data were used for training, and 10% was used for validation. The models updated the weights with a batch size of 32.

### Performance optimization with stochastic hill climbing

For the optimization of metabolite quantifications in plasma analysis, we implemented the stochastic hill-climbing algorithm with the output from CRNN as the starting point^30^. More specifically, CRNN can predict metabolites’ concentrations within a decent range, which could reconstruct a portion of the original plasma NMR spectrum (Fig. 2c). To further optimize the concentration values, we first globally correct the positional variations for all the metabolites. For each reconstructed spectrum, we shifted it within the region from -150 to 150 steps, with the negative sign denoting the direction of smaller chemical shift values. Under each shifted scenario, we evenly split the reconstructed and original spectra into 30 regions. We accumulated the regional correlations between the two spectra. The shift step, which raises the highest correlation score, will be used for global positional correction for the reference library (Fig. 2d).

With positions better aligned, we used stochastic hill climbing to iteratively fine-tune the mapping results for each component (metabolite, anticoagulant, lipoprotein) within the plasma^30^. In detail, the output from CRNN with global positional correction is taken as the starting point (Fig. 3a). In each iteration, we optimized one component at every step by multiplying its concentration randomly from 0.2 to 10 and shifting its positions randomly from -50 to 50 steps, with the negative sign denoting the direction of smaller chemical shift values. After the adjustment, we reconstructed the spectrum again based on the updated values. Mean squared error is calculated between the reconstructed and the original spectra. If mean squared error decreases compared to the previous value, we will keep the metabolite’s updated position and concentration. Otherwise, we will preserve the initial values. After 500 iterations, the mean squared error converged with no improvement (Supplementary Fig. 5). Users can also set the parameters to optimize only on position or concentration, further boosting the quantification performance.

### Evaluation for post-quantification results

With the estimated relative concentrations and the position-adjusted library from NMRQNet, we can reconstruct the whole plasma spectrum to compare with the original spectrum (Fig. 3c). The overlapping conditions could be evaluated using root mean squared error as the metric (Fig. 5a). We can also calculate the pixel-to-pixel residuals between the two spectra and check the residual plot (Fig. 5c). In addition, at the metabolite level, we can reconstruct metabolite-specific clusters and align them with the plasma spectrum to visually inspect the quantification performance (Fig. 4c, e). With manual cluster annotation or quantification results from the NMR expert available, we can further compare the estimated area under the curve or relative concentrations with the ground truth (Fig. 4a).

### Parameter settings for metabolites quantification methods

#### rDolphin

We followed the instructions on the GitHub repository of rDolphin: https://github.com/danielcanueto/rDolphin/blob/master/vignettes/rDolphin_introduction.Rmd.

The library is the same library rDolphin used for blood sample analysis in their paper. The parameter *Chemical shift tolerance (ppm)* in the ROI profile is tuned from 0.002 (default) to 0.02 to optimize the mapping results.

*Batman*. We followed the instructions of the *InstallAndTest*.*pdf* file on the R-project repository of Batman: https://r-forge.r-project.org/scm/viewvc.php/documentation%20and%20test/?root=batman. We extracted the overlapping 31 metabolites from Batman internal profiles as the library. We used all the default parameters in the MCMC model for 39 out of 40 samples. For one sample, *nItBurnin* is set to 3000 (default is 4000) to eliminate the reported error while running *batman* function. Batman takes a much longer time compared to other methods. It took around 6.5 hours to analyze one spectrum.

#### ASICS

We followed the instructions on ASICS’s Bioconductor repository: https://bioconductor.org/packages/release/bioc/vignettes/ASICS/inst/doc/ASICSUsersGuide.html. The library is customized with the library of NMRQNet for plasma analysis. The *max*.*shift* parameter is tuned from 0.02 (default) to 0.05, 0.08, and 0.1.

### Simulation of the training set for nine metabolites

We used nine metabolites first to test the feasibility of the proposed NMRQNet workflow. The nine metabolites include choline, glycine, myo inositol, cysteine, glutamate, glucose, lysine, tryptophan, and leucine. These nine metabolites were representatives of metabolites with different pattern complexities and chemical shift values. The number of metabolites and metabolite candidates are randomly determined in simulated samples. Since the background noise and potential contamination in real NMR spectra are hard to be captured by the simulated data, we manually added these noises by generating both the simulated and real pure spectra for every metabolite. Moreover, during the simulation process, for each simulated mixture, we randomly chose one metabolite and mixed its real NMR spectrum with the rest of the simulated spectra from other metabolites. It can help embed actual variations within simulated samples and prevent them from being over-complicated by adding too much background noise.

### Simulation of the training set for plasma samples

The plasma NMR spectrum has signals raised by multiple chemical compounds, including dominant metabolites, lipoprotein clusters, calibration standards, and anticoagulants. To include all the potential components, the real DSS (calibration standard) and different anticoagulant spectra were experimentally acquired. All potential lipoprotein clusters are annotated by a chemist and fitted with normal distributions. Peak signatures for 38 dominant metabolites were extracted from the reference library. To simulate the pattern variations, we scaled peak widths to generate dynamic metabolite clusters. Lorentzian-curve function further transformed the peak signatures into the metabolite spectra^31^. For positional variations, the metabolite spectra shifted within the neighboring region at the maximum step of 0.03 ppm. Spectra within the region of 0.7 ppm to 4.1 ppm are preserved for explorations since it includes most metabolomics signals. All the simulated or experimentally acquired spectra for different components were randomly mixed to generate the training samples (Supplementary Fig. 1).

### Concentration curve fitting

In most biofluid samples and tissues, only a few metabolites have dominant concentrations, and most metabolites have low concentrations. To simulate the spectra with metabolite concentrations distributed as in the real samples, we learned the distributions from real annotation results. Seven samples were prepared, including organoids, human plasma, cerebellum, cell, amniotic fluid, fly brain, and cell media. An NMR expert used Chenomx manually identify and quantify the metabolite concentrations within different samples. We fit the quantification results with a gamma distribution for every sample (Supplementary Fig. 1f). Then, in the simulation, concentrations are randomly sampled from the estimated gamma distributions. In the nine-metabolite study, we randomly used the seven fitted gamma distributions to generate concentrations. While in the plasma study, we used the fitted curve from the human plasma sample only to generate concentrations.

### Synthesize lipoprotein and EDTA clusters

Plasma is a complicated mixture with not only metabolites but also a large number of lipoproteins. Since lipoproteins usually have dominant clusters, learning their patterns will benefit the quantification of metabolites within the plasma^36^. Taking one plasma spectrum as a reference, the NMR expert identified the positions for lipoproteins clusters, which is further confirmed by previous lipoprotein analysis^36^. Then, computationally, we extracted these lipoprotein clusters and fit them with Lorentzian and normal functions. Overall, normal distributions have better fitting performance. Thus, we selected the normal-distribution curve with the smallest residual for each cluster as the simulated lipoprotein cluster within our library (Supplementary Fig. 1d). Finally, we simulate 24 lipoprotein clusters using normal functions.

In the simulated training set, we used the experimentally generated EDTA spectrum to represent EDTA features within the mixture, which only have free EDTA peaks. However, in plasma spectra, there are extra dominant clusters raised by EDTA’s calcium and magnesium complexes^34^. Thus, in the quantification process, mapping these complexes to prevent the misalignments of metabolite clusters is also necessary. To reconstruct these clusters, the NMR expert annotated their positions from one plasma spectrum. Together with two free EDTA peaks, two clusters from the EDTA-calcium complexes, and one singlet from the EDTA-magnesium complexes, we used Lorentzian curves to fit and simulate these clusters. The simulated clusters from free EDTA and its complexes are also added to the library during the quantification and optimization process. Their initial value is taken from CRNN’s EDTA prediction.

### NMR spectra preprocessing

For each raw ^1^H NMR spectrum, interference from the machine or other sources may raise distortions within the spectral patterns. And these distortions and alterations may result in bias in identifying or quantifying metabolites within the spectra. Martin et al. developed an NMR spectrum preprocessing pipeline implemented in PepsNMR, an R package^45^. The baseline correction method in PepsNMR was applied to estimate and remove the baseline from the mixture to reach a smooth background line (Supplementary Fig. 13a). For a general NMR spectrum, the informative metabolite region is from -1 to 11 ppm on the chemical shift axis. For nine metabolites, the whole region was kept for study. While in plasma analysis, most metabolites have their spectral signals clustered within the region of 0.7 to 4.1 ppm. The spectra within this region are preserved for later analysis to ensure spectral abundance and computational efficiency. An interpolation step was further applied to scale the NMR spectra to fit the input of the CRNN model. Finally, to ensure all the mixtures are identified and quantified on the same scale, normalization methods are mandatorily applied to eliminate the variations raised from samples or the system. The identification model used a min-max normalization to scale the spectrum from 0 to 1 (Supplementary Fig. 13b). While in the quantification model, the spectra were normalized to have areas under the curve to a constant sum, 1 (Supplementary Fig. 13c).

### Data preparation

In the nine-metabolite study, mixtures with known compositions include 0.5mM Glucose, 2.2mM Glycine, 0.07 mM Choline, 6.8mM Glutamic acid, 0.3mM Myo-inositol, 1.4mM Cysteine. The solution was prepared in pH 7.0 1XPBS as a mixture with 0.5 mM DSS as the internal standard. The final solution was 500uL with 10% D2O and 90% H2O and carefully transferred to the 5mm NMR tubes. ^1^H NMR measurements of the standard samples were performed using the 800MHz Bruker Avance instrument. The spectra were acquired using the avance-version noesypr1d pulse sequence with pre-saturation during relaxation delay and mixing time. The temperature was adjusted before the experiment. The acquisition mode was DQD with a size of fid 65K. The dummy scans were four, and the number of scans equaled 128. The spectral width was 20.56 ppm, and the acquisition time was 1.99 seconds. The Fid resolution was 0.5Hz, and the filter width was 4 million Hz. The receiver gain was 50, and the dwell time was 30.4 usec.

In plasma analysis, the 40 human plasma samples from the Mecp2 duplication syndrome families were collected with EDTA as the anticoagulant. The plasma samples were prepared to generate NMR spectra with the following steps. The 450uL human plasma was mixed with 50uL 5mM DSS as the internal standard. The final solution was 500uL with 10% D2O and 90% H2O and carefully transferred to the 5mm NMR tubes. ^1^H NMR measurements of the standard samples were performed using the 800MHz Bruker Avance instrument. The 1D spectra were acquired using the avance-version noesypr1d pulse sequence with pre-saturation during relaxation delay and mixing time. The 2D TOCSY spectra were acquired using avance-version dipsi2esgpph, the homonuclear Hartman-Hahn transfer using DIPSI2 sequence for mixing and phase sensitive, as well as water suppression using excitation sculpting with gradients. The acquisition mode is DQD, and the sizes of fid were 4k for F2 and 2k for F1, respectively. The dummy scans were 160, and the number of scans was 24. The loop count for td0 was 1. The spectral width was 13.95ppm for both F2 and F1. The receiver gain was 203.

For mouse plasma samples, with heparin and citrate separately as anticoagulants, 300 uL mouse plasma samples were prepared in 150 uL 1XPBS as a mixture, with 50uL 5mM DSS as the internal standard. The final solutions were transferred to the 5mm NMR tubes. ^1^H NMR measurements of the standard samples were performed using the 800MHz Bruker Avance instrument. The spectra were acquired using the avance-version noesypr1d pulse sequence with pre-saturation during relaxation delay and mixing time. The temperature was adjusted before the experiment. The acquisition mode was DQD with a size of fid 65K. The dummy scans were 16, and the number of scans equaled 2K. The spectral width was 20.56 ppm, and the acquisition time was 1.99 seconds. The Fid resolution was 0.5 Hz, and the filter width was 4 million Hz. The receiver gain was 203, and the dwell time was 30.4 usec.

## Data availability

The data synthesized and generated for this study is available at https://github.com/LiuzLab/NMRQNet

## Code availability

NMRQNet is implemented with R and Python. The codes and tutorials for NMRQNet are available at https://github.com/LiuzLab/NMRQNet

## Acknowledgement

We thank Cemal Karakas, MD, Soumya Patti, PhD, Marina Vannucci, PhD, and Christine Peterson, PhD for their helpful comments and inputs for this work; Advanced Technology Core support of NMR and Drug Metabolism core, the National Institute of General Medical Sciences (5R01GM120033 to M.M.S.). the Eunice Kennedy Shriver National Institute of Child Health & Human Development of the National Institutes of Health under Award Number P50HD103555 (M.M.S) for support of the Bioinformatics Core and the NMR and Drug Metabolism Core. The content is solely the responsibility of the authors and dose not necessarily represent the official views of the National Institutes of Health. The work was partially supported by the Cynthia and Antony Petrello Endowment (M.M.S.), the Chao Endowment (Z.L.) and the Huffington Foundation (Z.L.).

## Author contributions

Z.L., M.M.S., and W.W. planned and initiated the project. Z.L., M.M.S. supervised the project. W.W. designed, set up the algorithm, and analyzed the data. L.M. performed the NMR experiments. W.W., L.M. wrote the manuscript.

## Corresponding authors

Correspondence to Zhandong Liu.

## Competing interests

The authors declare no competing interests.

## Notes

### Competing Interest Statement

The authors have declared no competing interest.

