## Supplementary Figures for "NMRQNet: a deep learning approach for automatic identification and quantification of metabolites using Nuclear Magnetic Resonance (NMR) in human plasma samples"

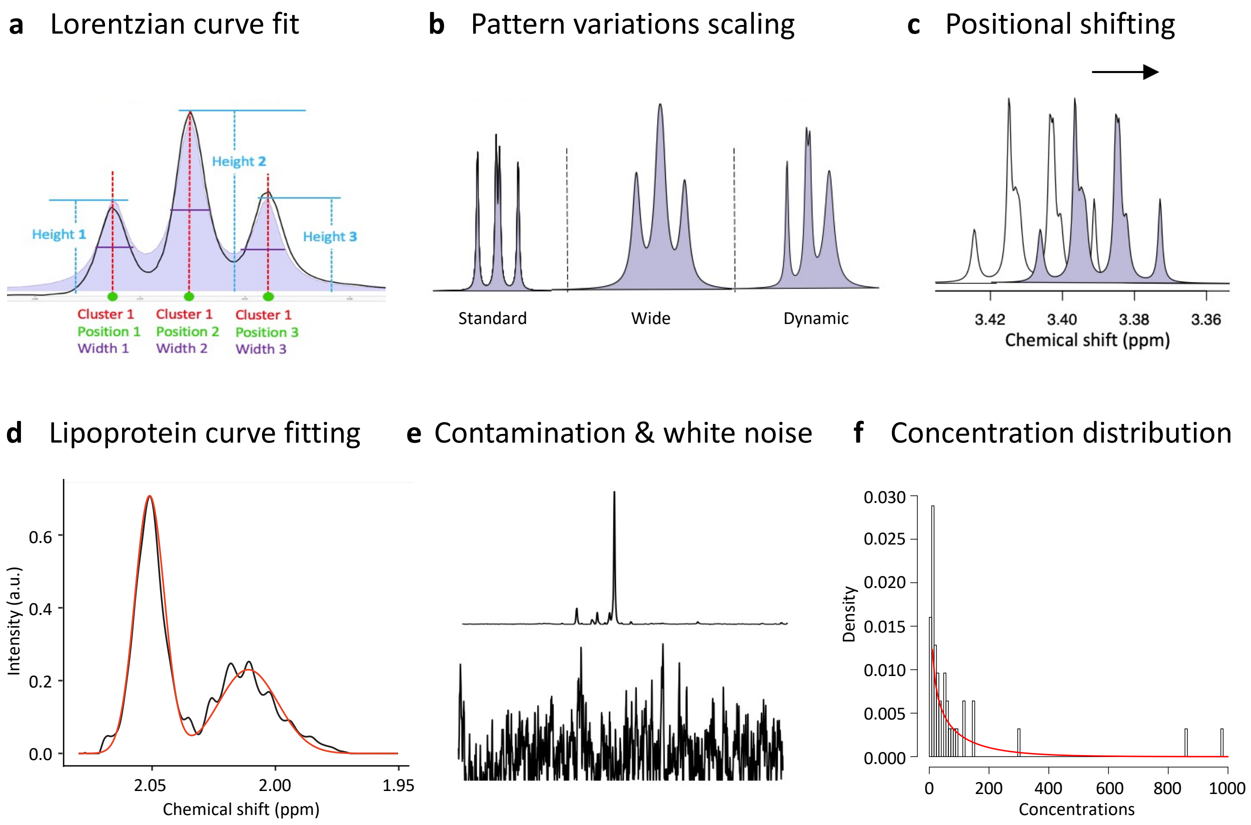


**Fig. S1**. Simulate training samples with variations and noise embedded. a, Lorentzian curve converts the peak signatures in the reference library into the whole NMR spectrum. b, Data augmentation is applied by altering the peak width scales during the Lorentzian-curve fitting process to simulate pattern variations for the same metabolite cluster. c, Data augmentation is applied by shifting the metabolite cluster within the neighboring regions to imitate positional variations within the real NMR spectra. d, Potential contamination signals, and background noise are added into the simulation process. e, Curve-fitting for real samples manual quantification results. In the simulation, concentrations within each mixture are generated based on the fitted curve.

**Fig. S2.** Identification results on known mixture under different temperatures. a, Positional variations under different temperatures. b, Heatmap for identification probabilities across temperatures.

**a**

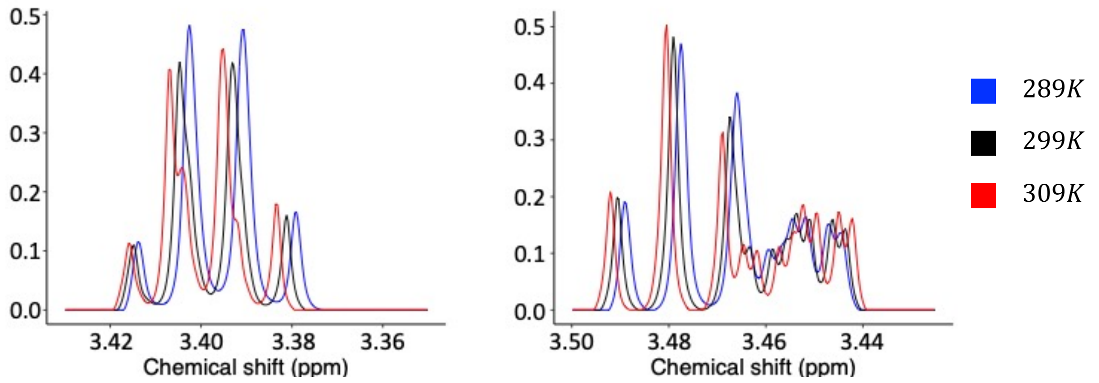


**b**

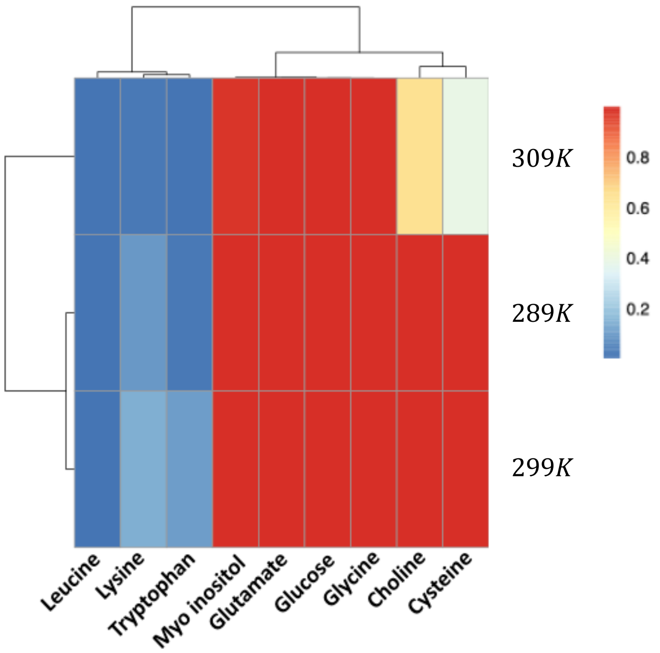

**Fig. S3.** Quantification results on a nine-metabolite simulated sample.

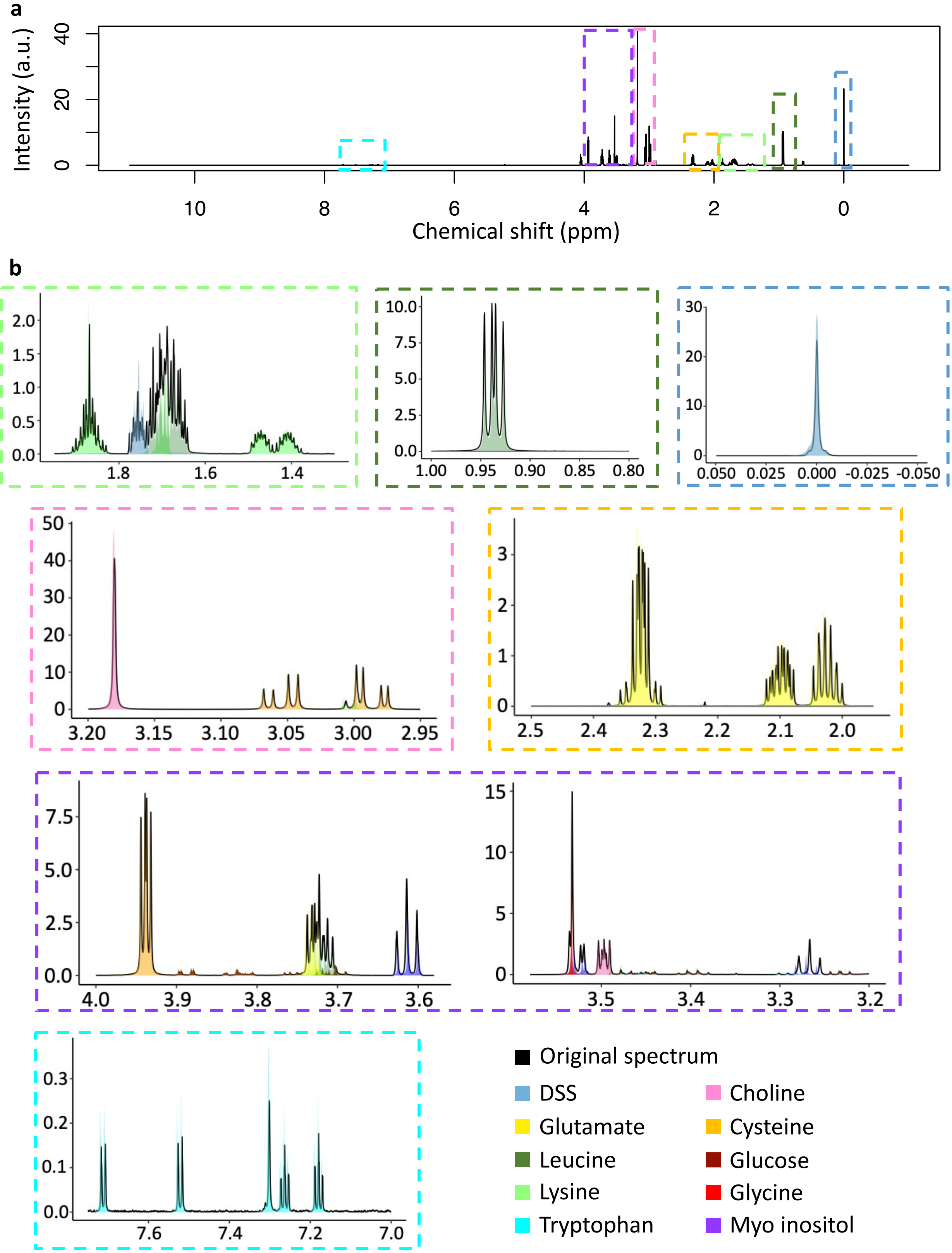

**Fig. S4.** Quantification results on a nine-metabolite known mixture under $299K$ .

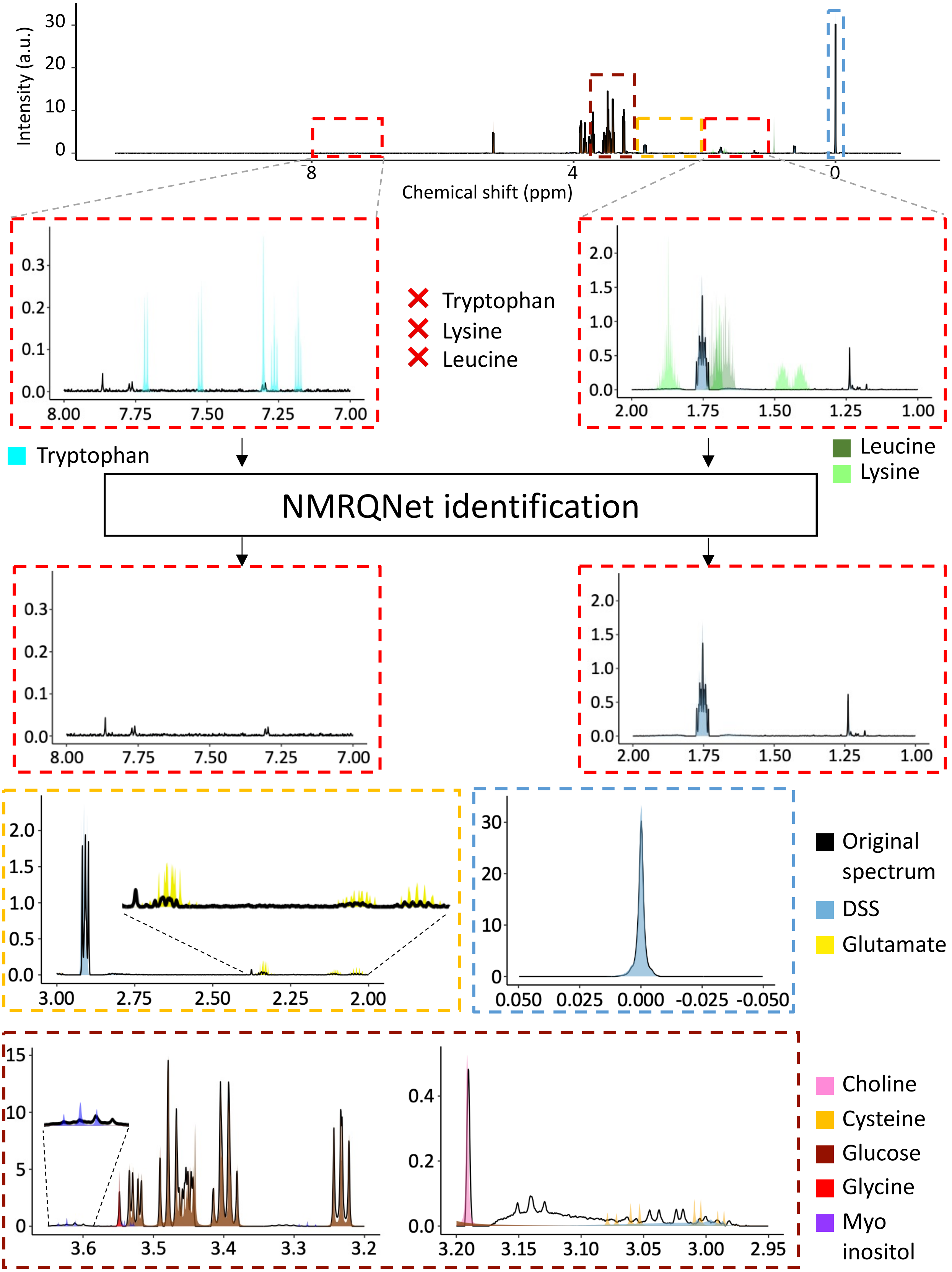

**Fig. S5.** Mean square error plot over the 500 iterations in local optimization process.

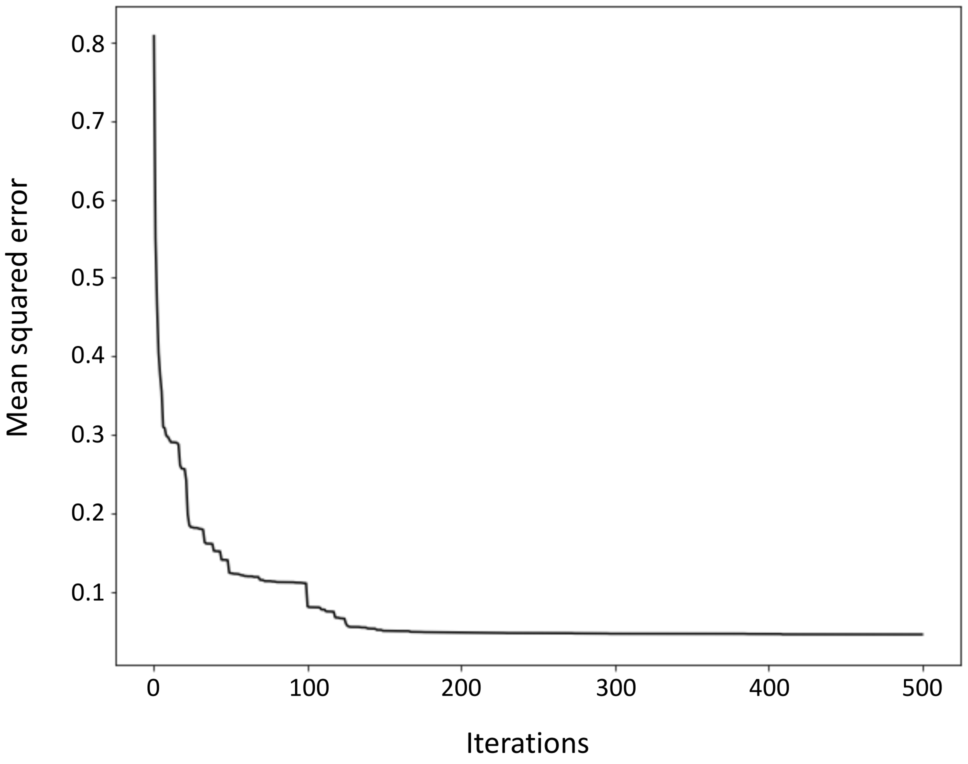

**Fig. S6.** Quantification results on plasma samples from different species with diversified anticoagulants. a, Mouse plasma with heparin as anticoagulant. b, Mouse plasma with citrate as anticoagulant. c, Human plasma with EDTA as anticoagulant.

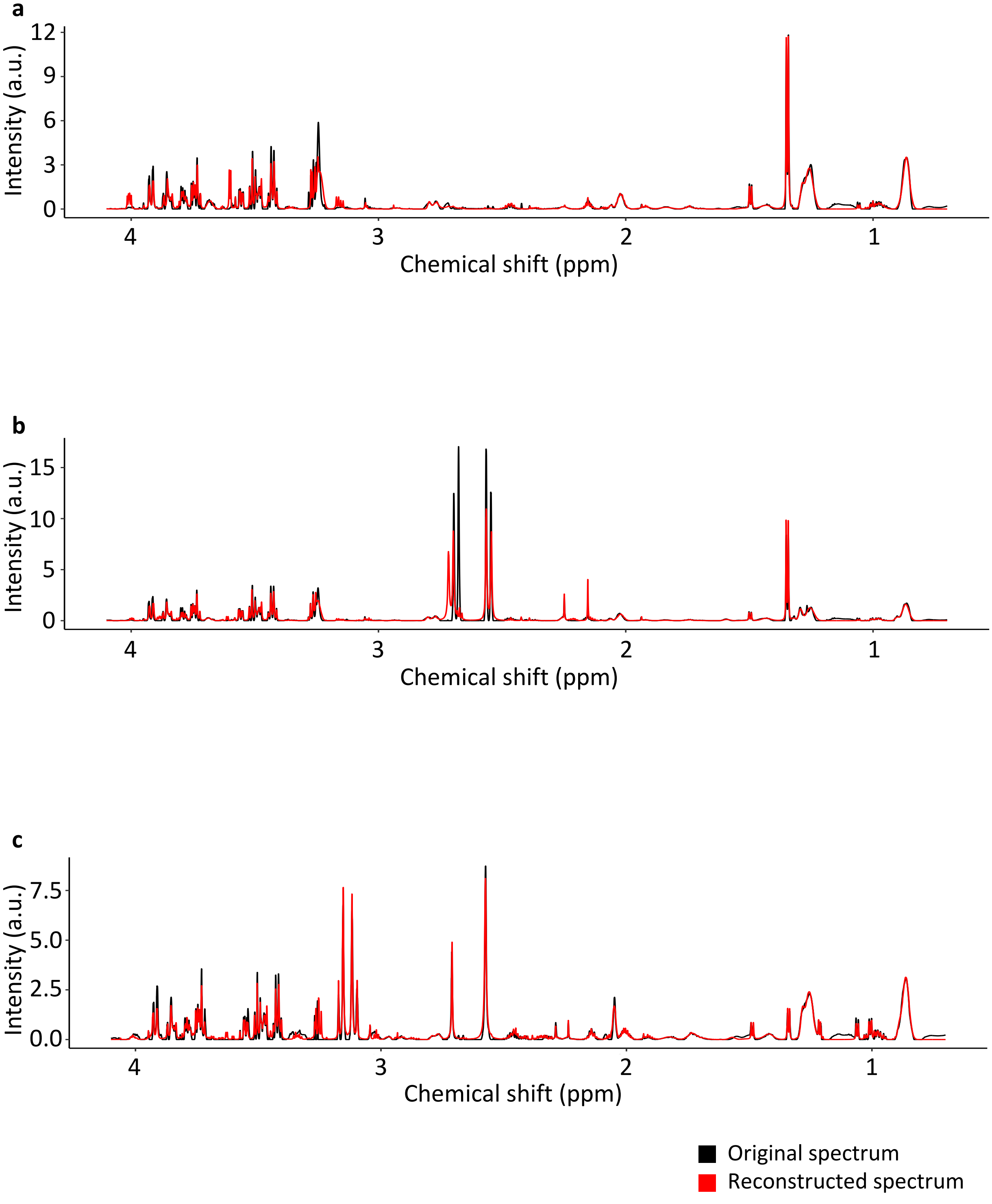

**Fig. S7.** ASICS mapping results for the same sample with different max.shift values

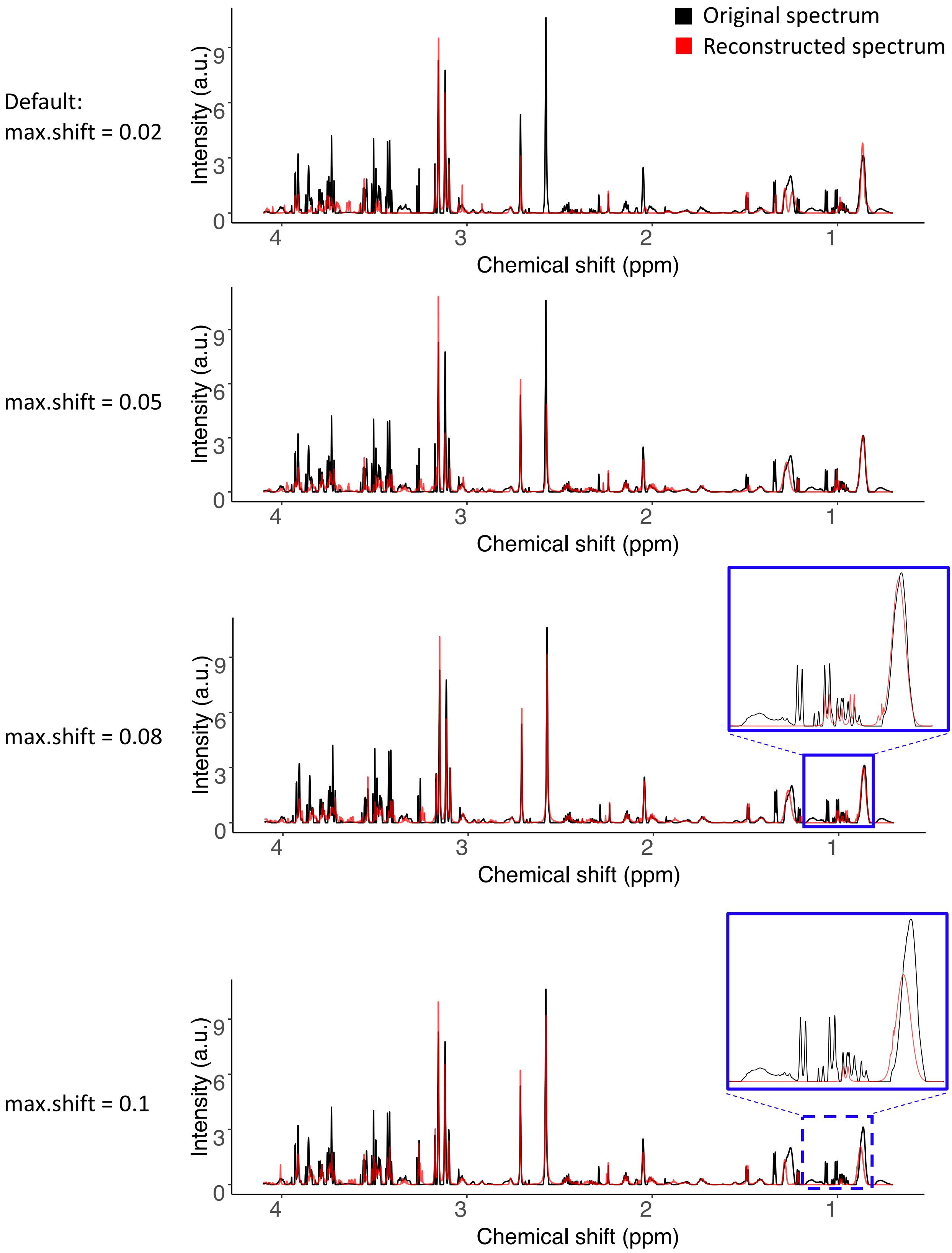

**Fig. S8.** In ASICS, optimal max.shift is not consistent across samples within in the same group

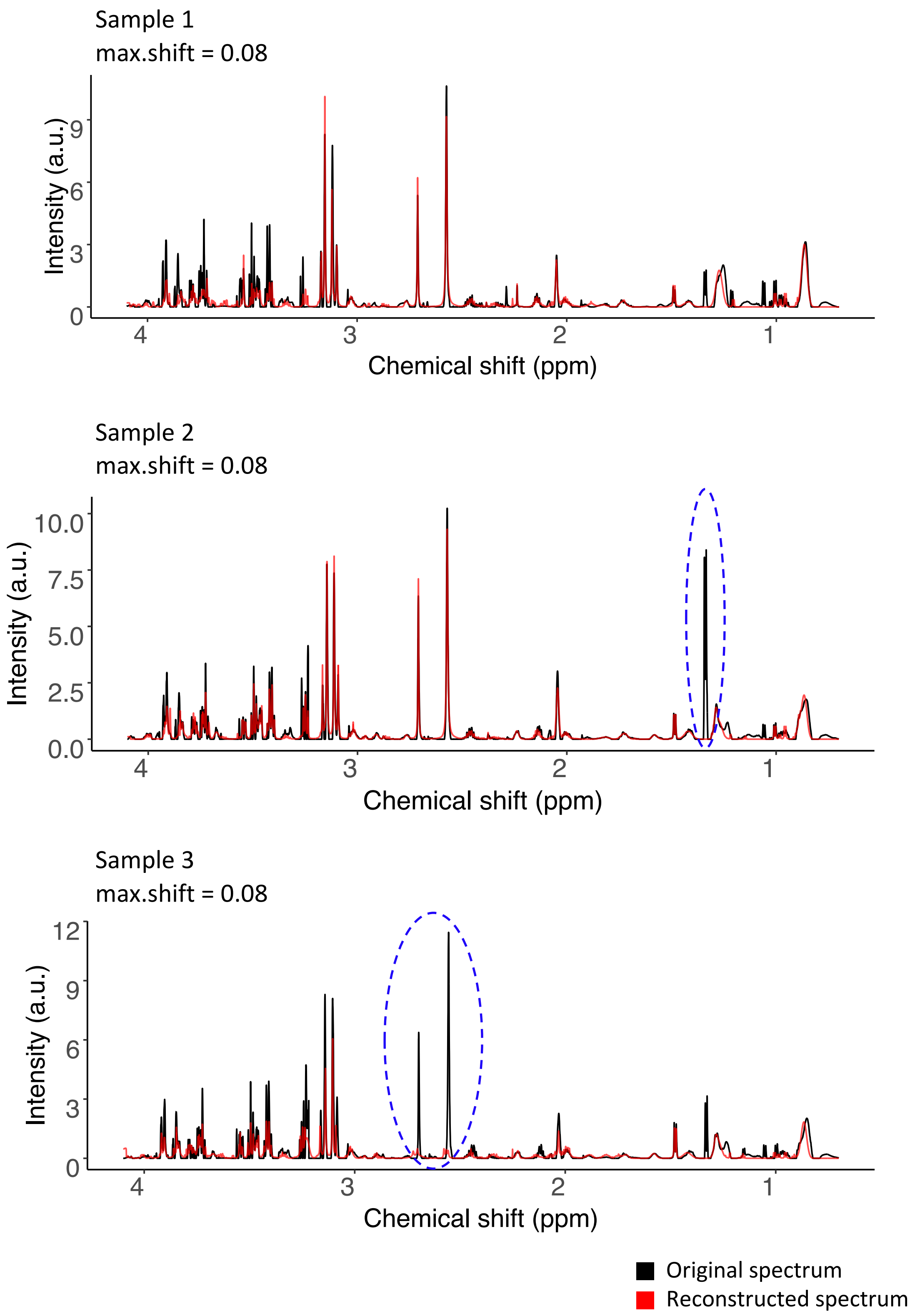

**Fig. S9.** Explore the sample where ASICS has lower RMSE compared to NMRQNet. a, Original NMR spectrum. b, Reconstructed residuals between NMRQNet and ASICS. c,  Regional reconstruction RMSE comparisons. D, Spectral signals within the region of 1.0 to 1.5 ppm. e, Residuals comparisons within the region of 1.0 to 1.5 ppm.

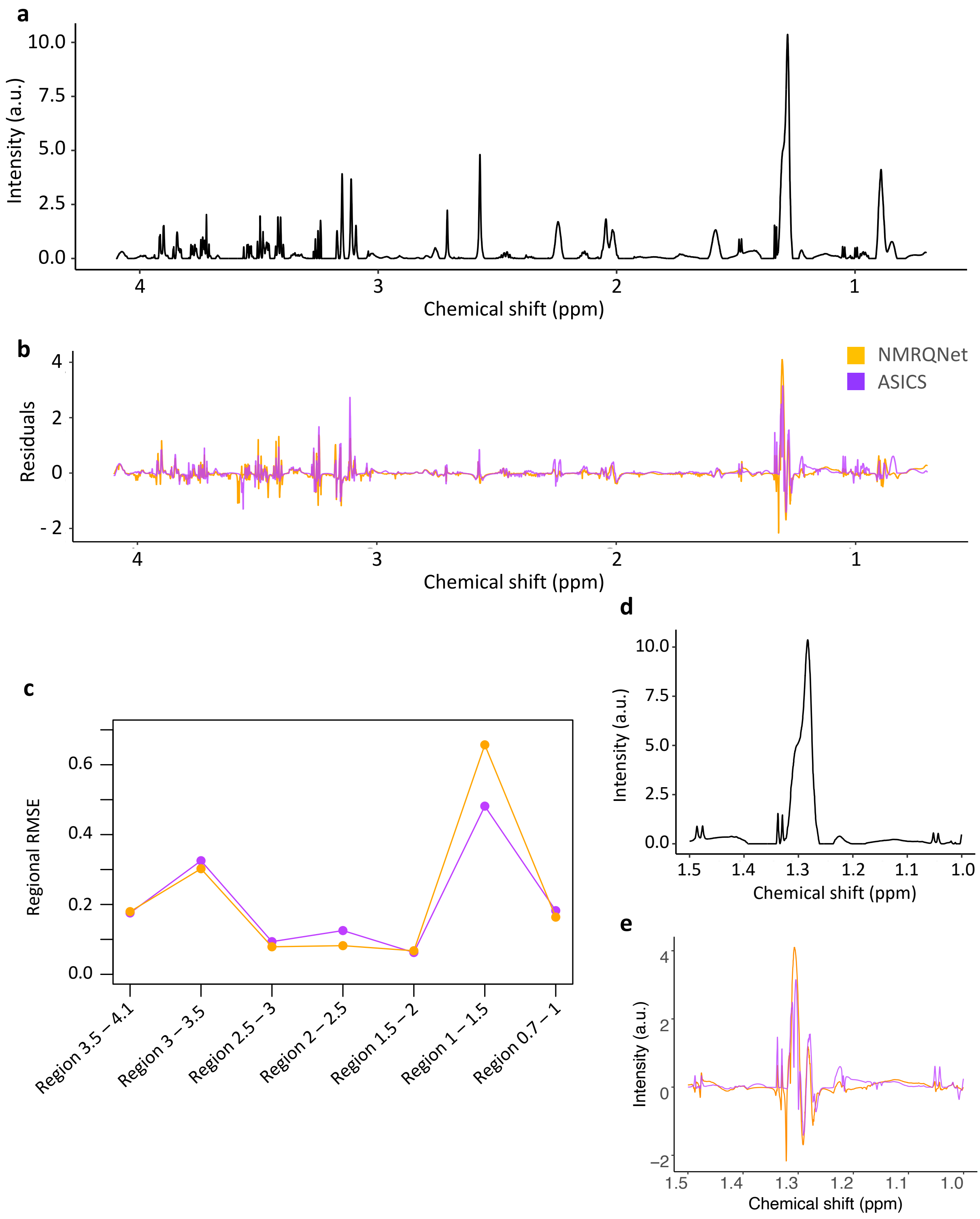

**Fig. S10.** PCA on 38-metabolite estimated concentrations for 40 samples from Mecp2 duplication syndrome families. a, PCA plot. b, PCA variable importance plot.

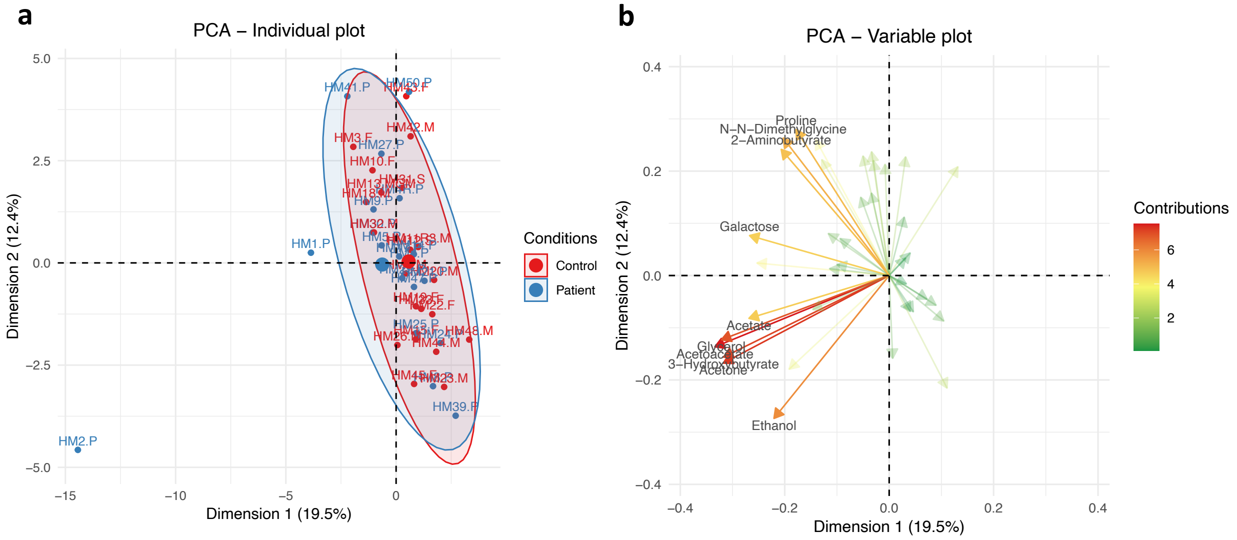

**Fig. S11.** PCA on 38-metabolite estimated concentrations after removing the outlier. a, PCA plot. b, PCA variable importance plot.

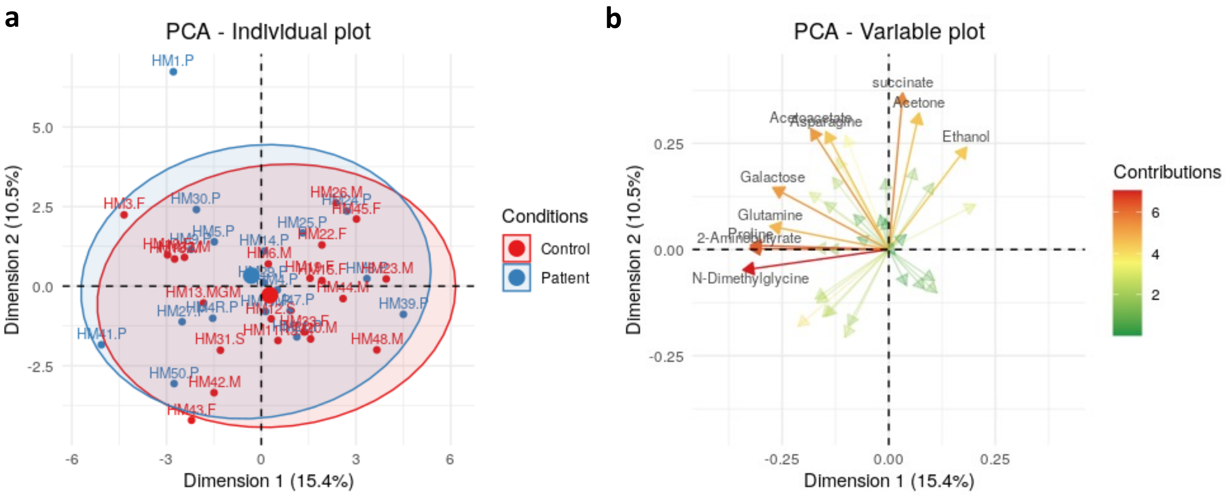

**Fig. S12.** Logistic regression on 38-metabolite estimated concentrations in the prediction of control and patient group. a, Leave-one-out predictions on 39 samples. b, Precision-recall curve. c, Estimated coefficients for different metabolites.

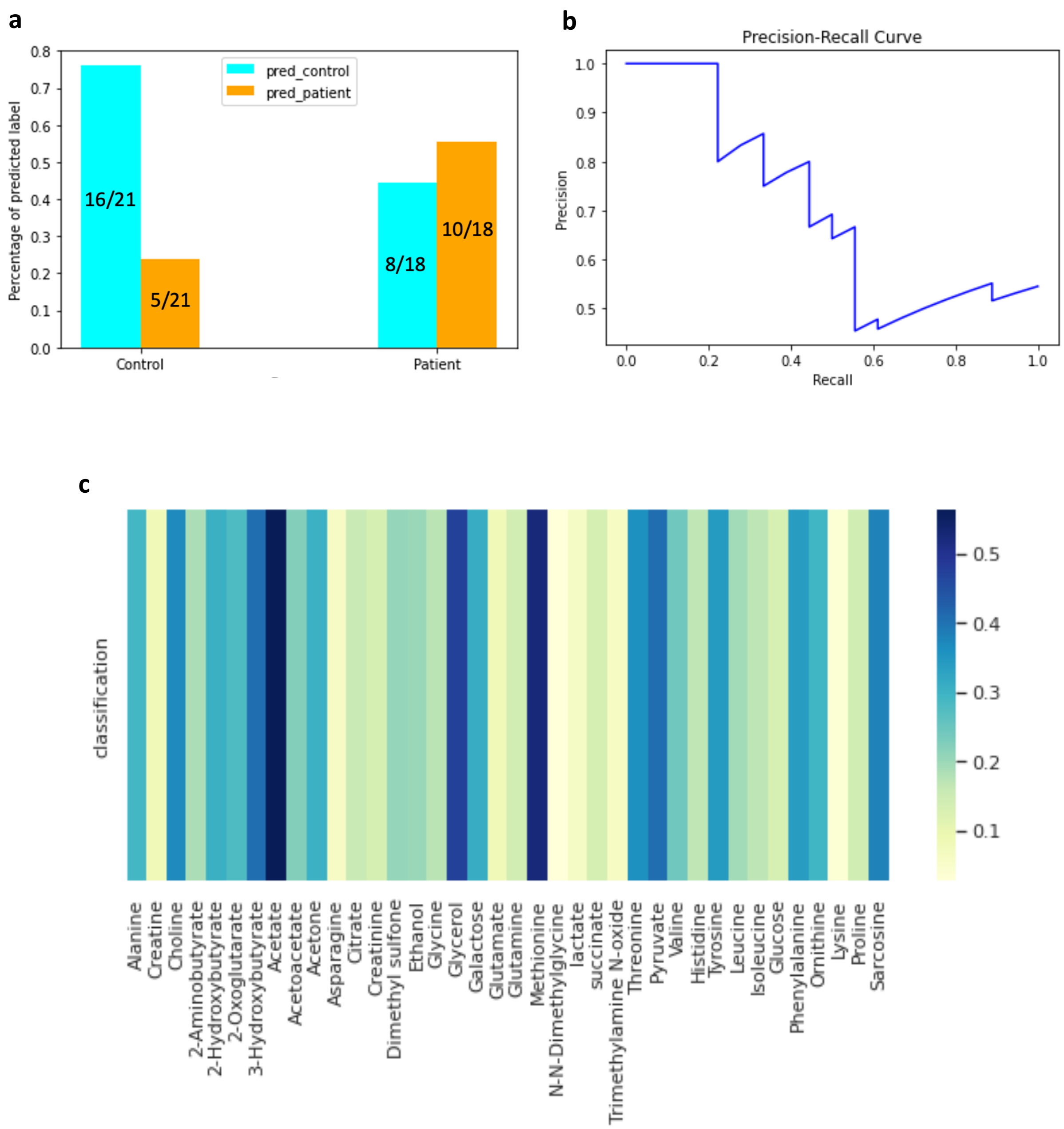

**Fig. S13.** Preprocessing steps for real NMR spectra. a, Baseline correction for raw plasma NMR spectra. b, Normalization. Maximum intensity value equal to 1. c, Normalization. Area under the curve equals to 1.

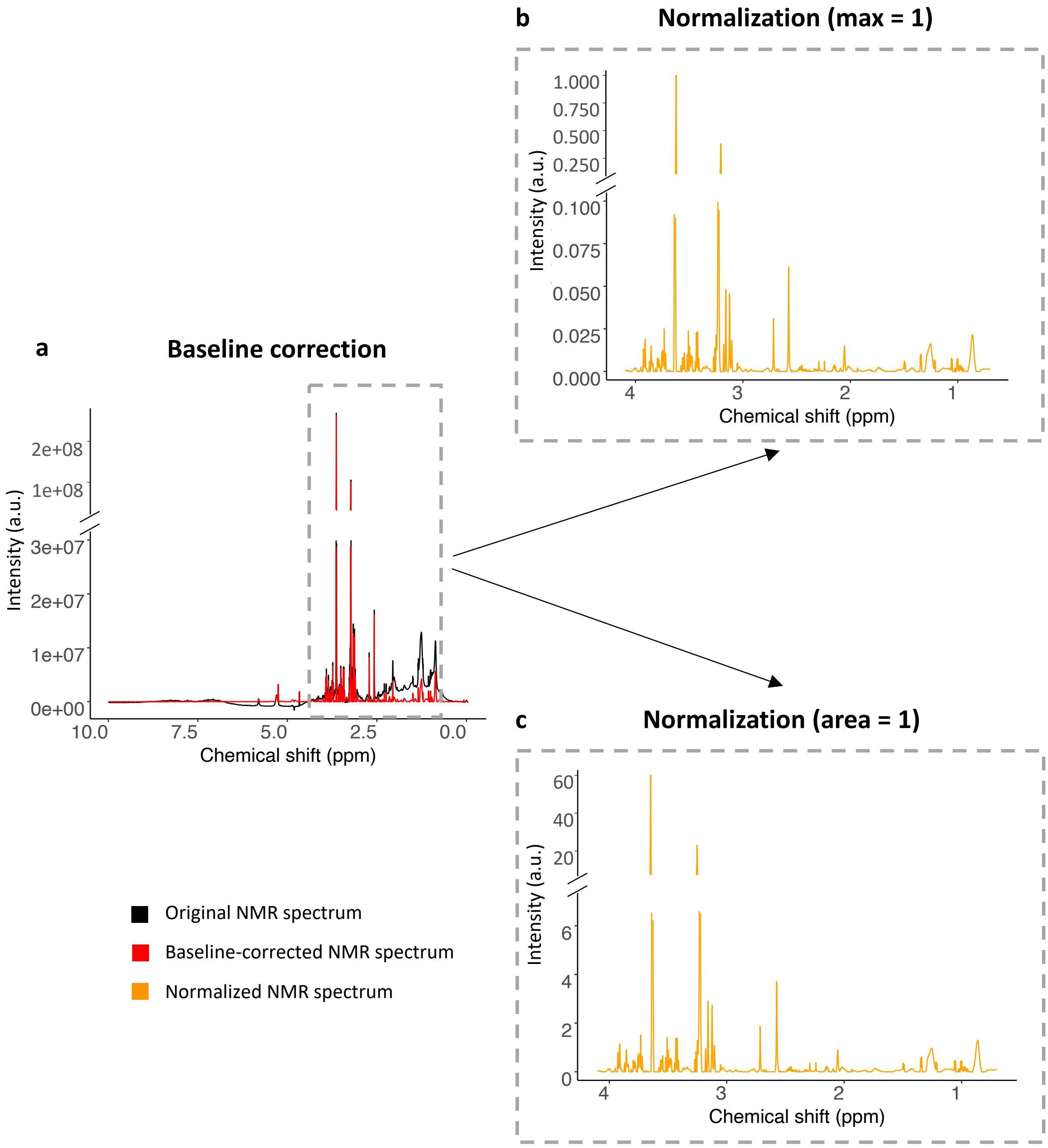

**Supplementary Table S1:** Sample-level identification accuracy for nine metabolites on simulated samples

| Accuracy | # Sample | Percentage |
|---|---|---|
| 9/9 | 6422 | 92.06% |
| 8/9 | 502 | 7.20% |
| 7/9 | 47 | 0.67% |
| 6/9 | 5 | 0.07% |
| < 6/9 | 0 | 0.00% |

**Supplementary Table S2:** Metabolite-level identification accuracy for nine metabolites on simulated samples

| Metabolite | Accuracy |
| --- | --- |
| Choline | 0.985 |
| Cysteine | 0.995 |
| Glucose | 0.990 |
| Glutamate | 0.993 |
| Glycine | 0.978 |
| Leucine | 0.994 |
| Lysine | 0.991 |
| Myo inositol | 0.991 |
| Tryptophan | 0.995 |

**Supplementary Table S3:** Experimentally curated known mixture from nine metabolites

| Metabolite | Concentration (mM) |
|---|---|
| Glucose | 5.50 |
| Glycine | 0.22 |
| Choline | 0.01 |
| Glutamate | 0.02 |
| Myo Inositol | 0.03 |
| Cysteine | 0.02 |
| Leucine | 0.00 |
| Lysine | 0.00 |
| Tryptophan | 0.00 |
